# Alternative splicing regulation in plants by effectors of symbiotic arbuscular mycorrhizal fungi

**DOI:** 10.1101/2023.09.20.558436

**Authors:** Ruben Betz, Sven Heidt, David Figueira-Galán, Thorsten Langner, Natalia Requena

## Abstract

Most plants in natural ecosystems live in association with AM (arbuscular mycorrhizal) fungi to survive under poor nutrient conditions and to cope with other abiotic and biotic stresses. To engage in symbiosis, AM fungi secrete effector molecules that, similar to pathogenic effectors, reprogram plant cells. Despite numerous effectors being predicted in the genome of AM fungi, only a few have been functionally characterized. Here we show that the SP7-like family, a Glomeromycotina-specific effector family, impacts on the alternative splicing program of their hosts. SP7-like effectors localize at nuclear condensates and interact with the plant mRNA processing machinery, most prominently with the splicing factor SR45 and the core splicing proteins U170K and U2AF35. Ectopic expression of two of these effectors in the crop plant potato changed the alternative splicing pattern of a specific subset of genes. Unravelling the communication mechanisms between symbiotic fungi and their host plants will help to identify targets to improve plant nutrition.

## Introduction

In nature, plants engage in multiple microbial interactions that ultimately determine plant health and productivity. During these associations, both plants and microbes exert all their capabilities to take control of the interaction, including the delivery of peptides, proteins and small RNAs to the other partner in order to manipulate their cellular program^1–4^. These effectors molecules are key components of the interaction between microbes and their host plants. For microbes, they allow them to remain invisible to the surveillance system of plant cells, to subvert defense reactions and/or to manipulate plant metabolism. Although most effectors characterized belong to pathogenic microbes, increasing evidence shows that beneficial associations also rely on the delivery of effector molecules to shape the mutualism^5,6^.

The increasing availability of sequenced genomes from different AM fungal species and prediction tools has revealed the existence of many putative effector proteins^7–11^. However, only a handful of them have been functionally characterized and shown to play a role in the symbiosis^12–16^. Microbial effectors have a wide range of possible final locations *in planta* ranging from the apoplastic space to different intracellular compartments including the nucleus. This allows effectors to find their specific targets and control multiple cellular processes^17–19^. Nuclear effectors have been shown to modulate hormonal pathways, host metabolism or immune responses by different mechanisms that include interactions with nuclear proteins, DNA or RNA^17,19^. Interestingly from the five effectors characterized in the fungus *Rhizophagus irregularis*, three are located in the nucleus^12,14,16^ This might reflect the large number of predicted nuclear effectors in this fungus, as revealed in an analysis of *R. irregularis* secretome during symbiosis, in which from ca. 300 putative effectors, almost one third had a predicted NLS^15^, pointing to the nucleus as a key target for arbuscular mycorrhizal symbiotic effectors. Interestingly, two of the few characterized effectors from rhizobia, ErnA and Bel2-5 are also targeted to the plant nucleus and shown to promote nodulation^5^. But also, numerous effectors of plant pathogens have the nucleus as their playground^17,18^.

From all nuclear targeted effectors characterized so far, only a few have been shown to impact on pre-mRNA splicing. This is remarkable because alternative splicing is a key regulatory mechanism of gene expression in plants^20–23^, where it is estimated that between 60-70% of intron containing genes undergo AS^24^. In the crop plant potato from *ca.*33.000 genes more than 44.000 transcripts are predicted, and 7000 genes are known to have more than one transcript (Phytozome v.13). Alternative splicing in plants contributes to the ability of plants to respond to environmental changes including abiotic and biotic stresses, but also to control developmental processes^20,22,25^. Furthermore, the role of AS in plant immunity is increasingly recognized with defense responses altered in mutants of splicing regulators^26–28^, altered splicing landscape during infection by pathogens^29–31^ or the discovery of effectors targeting AS^31–34^.

Pre-mRNA splicing occurs at the spliceosome, where besides the large ribonucleoprotein complex (U1, U2, U4, U5 and U6) many other accessory proteins participate. Among those, the SR proteins are a family of RNA-binding proteins with a prominent role in constitutive and in alternative splicing. They were first identified as chromatin interacting proteins, and have been shown to participate not only in pre-mRNA splicing but also in mRNA nuclear export and translation control^35,36^. A loss of function of the SR protein SR45 led to the surprising finding that this protein, besides its already known roles in development and abiotic stress^37–39^, was a suppressor of innate immunity in Arabidopsis^26^. SR45 and its human counterpart RNPS1 are members of the conserved Transformer-2 subfamily within the SR superfamily and are unusual splicing factors containing two arginine-serine (RS) rich domains flanking an RNA recognition motif (RRM) ^38^. While the RRM motif determines RNA-binding specificity, the RS domains participate in protein-protein interactions^40^. RNPS1 is a peripheral protein of the exon-junction-complex (EJC) that serves as platform for splicing regulatory complexes to maintain transcriptome surveillance by regulating alternative splicing and non-sense mediated decay^41,42^. In *Arabidopsis*, SR45 was also postulated as part of the EJC^43^ and shown to colocalize with U2AF^35^b, Cactin and other SR proteins in nuclear condensates^40,44^. Arabidopsis SR45 is even able to complement splicing of some pre-mRNAs in splicing-deficient HeLa cell S100 extract^37^.

Here we demonstrate that beneficial symbiotic AM fungi employ an effector protein family, widely conserved across the Glomeromycotina, that interacts with components of the plant alternative splicing machinery, most significantly with SR45, and modify the alternative splicing of a subset of genes when expressed *in planta*.

## Results

### The SP7-like effector family

SP7 belongs to a small family of proteins in the symbiotic arbuscular mycorrhizal fungus *R. irregularis* that shares a conserved modular structure, comprising a signal peptide, a series of hydrophilic imperfect repeats and, in several cases, a nuclear localization domain (Fig. 1a). Based on the previously identified sequences^7,8,12^, we searched on the new annotation of the *R. irregularis* genome^9,10^ and identified five *core* RiSP7-like sequences, RiSP7, RiSP2, RiSP5, RiSP31 and RiSP1 (Fig. 1a). Surprisingly, for four of them there are several highly conserved paralogues scattered through the genome in different chromosomes, with the exception of *RiSP2*. Only in two cases, more than one *SP7-like* gene is found in the same chromosome (Fig. 1b). Paralogues encode proteins that share between 85% and 96% amino acid identity (Supplementary Fig. 1), while the identity among the five core RiSP7-like proteins is no more than 49% (Supplementary Fig. 1). Because the AM symbiosis is formed by many different fungal species, we investigated whether similar effectors would exist in other AM fungi (Supplementary Tables 1 and 2). Amplification of two putative homologues from *Gigaspora margarita* and *Gigaspora rosea*^45,46^ and alignment of the proteins with the SP7-like sequences from *R. irregularis* revealed a similar structure and the presence of a characteristic signature that defines the family (Fig. 1a, Supplementary Figs. 1 and 2). Furthermore, a conserved intron preceding the first repeat was found in all SP7-like genes (Supplementary Fig. 2) including also other species such as *R. diaphanus* and *R. clarus*, suggesting a common gene ancestor for this effector family despite their sequence divergence (Fig. 1c). The exon-intron structure has been used to identify a common origin for other effectors, including the Y/F/WxC effector family from the pathogenic fungus *Blumeria graminis* f.sp. *hordei*^47^ or the Bicycle effectors from aphids^48^. Using the criteria of this conserved signature and the conserved intron, we found that SP7-like proteins are present in the Glomerales and Diversisporales, but there are so far no hints of them in more ancient orders such as Paraglomerales or Archaeosporales nor in other subphyla from the Mucoromycota such as Morteriellomycotina or Mucoromycotina (Fig. 1d and Supplementary Tables 1 and 2). Taken together, the conservation of the SP7-like family within the Glomeromycotina and its expansion in *R. irregularis* suggests that they could play an important role in the AM symbiosis.

**Figure 1.**
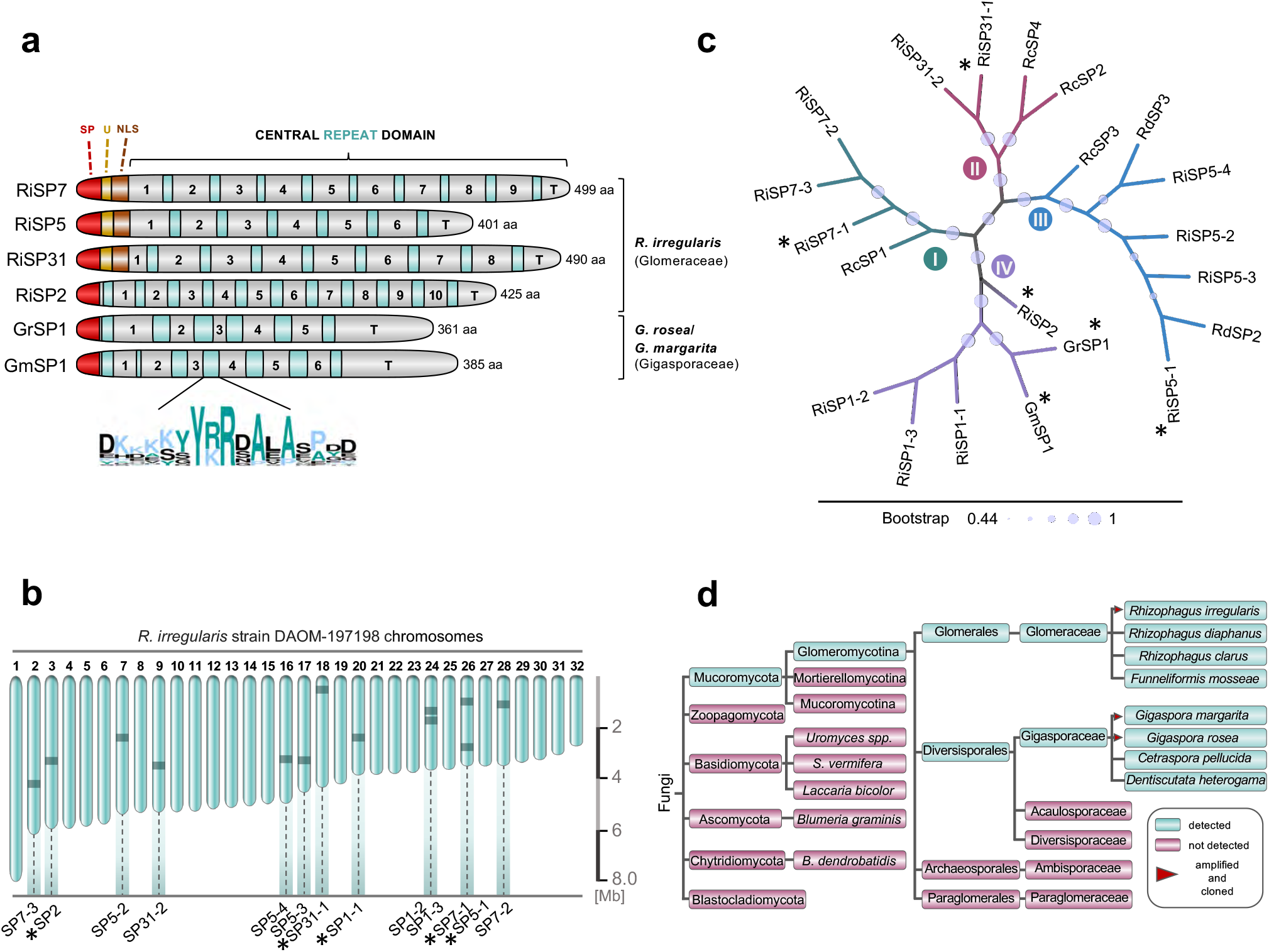
The multi-membered SP7 effector family is conserved and specific to AMF. **a** Protein features of cloned SP effectors. All effectors possess a signal peptide (SP, red, SignalP-5.0). For some members this is followed by a conserved 10-12 aa stretch of unkown function (U, yellow) and a consecutive bipartite NLS (brown, PSORTII). The repeat domain contains 6-11 imperfect hydrophilic repeat units. Variable aa (insertions/deletions/substitutions) define each repeat-subunits. A characteristic highly conserved island marks the end of each repeat unit (green coloured boxes, consensus logo). Terminal repeats (T) lack these islands. **b** Distribution of SP7 family effectors in the genome of *R. irregularis*. Chromosomes 1-32 of *R. irregularis* strain DAOM-197198 are depicted according to Manley *et al*., 2023. Relative positions of SP7 family members are indicated by dark bars. *R. irregularis* contains four core members (*SP7-1*, *SP1-1*, *SP2*, *SP5-1* and *SP31-1,* asterisks) that exist as paralogs in varying numbers. **c** Phylogenetic relationship of full length SP7 family members identified for different AMF species. Shown is an unrooted Maximum Likelihood tree (MEGA11, JTT modelling) of SP7 family members after ClustalW alignment. Bootstrap values (1000 replicates) are shown by branch circles. SP7-like effectors cluster into four distinct clades (I-IV). Tree was visualized using iTOL. Asterisks mark cloned effector members. Ri (*Rhizophagus irregularis*), Rc (*Rhizophagus clarus*), Rd (*Rhizophagus diaphanus*), Gm (*Gigaspora margarita*), Gr (*Gigaspora rosea*). **d** Analyses of genome and RNAseq data (NCBI Sequence Read Archive (SRA)) using SP7 family effector sequences as query revealed the ubiquitous distribution of SP effector genes or transcripts in the subphylum Glomeromycotina, while they were not found in any other of the analyzed fungal taxa (see Supplementary Table 1 and 2 for details). Red triangles mark species that were used to clone full length genomic or cDNA effector sequences.

### SP7-like proteins localize to the plant nucleus in nuclear condensates

We had shown previously that RiSP7 localizes in the nucleus of the plant when expressed *in planta* as predicted by the presence of an NLS signal^12^. However, several of the SP7-like members of the gene family, including those from *Gigaspora* species, apparently lack a clear NLS. Therefore, in order to investigate their localization, we expressed several of them ectopically in *Nicotiana benthamiana* epidermal cells, fused to GFP either N-terminally or C-terminally. Surprisingly, all of them showed predominantly nuclear localization despite lacking a predicted NLS (Fig. 2a). An exception was RiSP31 that only localized to the nucleus when carboxy-terminally labelled, albeit in a very low amount, despite possessing a distinct NLS. All analyzed SP7-like effectors were also present in the cytoplasm to a certain extent, with the exception of RiSP2 that was solely located in the nucleus (Fig. 2b). Furthermore, it could be shown that in some cases RiSP7 and RiSP5 localized to P (processing bodies) in the cytoplasm (Supplementary Fig. 3). A higher magnification of the nucleus showed that often SP7-like proteins accumulated in nuclear bodies in inhomogeneous numbers and sizes, but never in the nucleolus (Fig. 2a and Supplementary Fig. 4). Different types of nuclear bodies have been identified forming as a result of phase separation processes, often involving the recruitment of intrinsically disordered region-containing proteins^18^. Phase separation is mediated by molecules with the ability to form many interactions allowing the formation of subcellular or intra-organelle membrane-less compartments termed biomolecular condensates^49,50^. From those, nuclear condensates have a recognized role as mRNA processing organelles. All SP7-like predicted proteins are almost entirely intrinsically disordered (Supplementary Fig. 5), suggesting the possibility that they might have multiple interaction partners and be components of biomolecular condensates.

**Figure 2.**
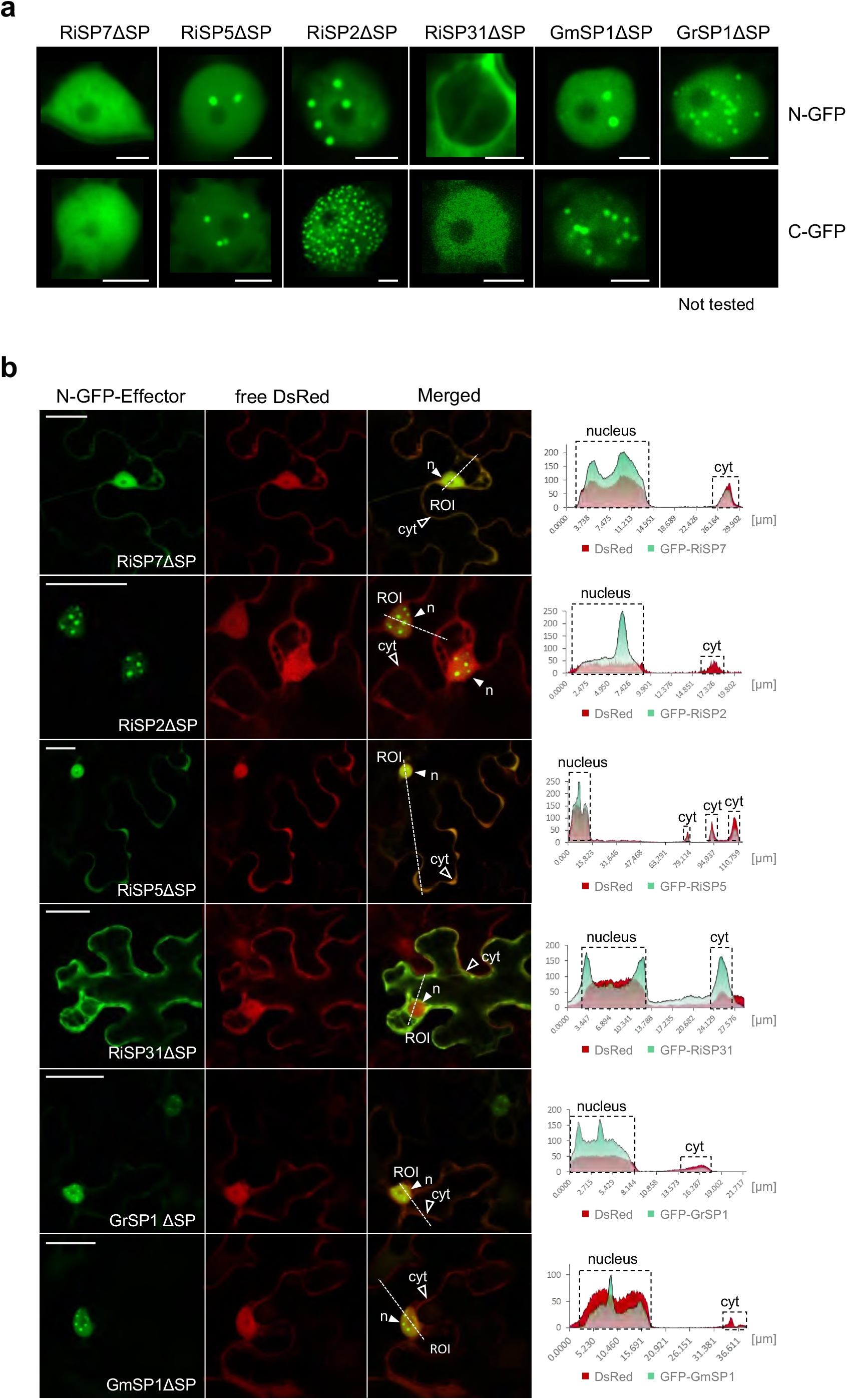
The SP7-like effector family localizes to the plant nucleus and accumulates in nuclear condensates. **a-b** Subcelluar localization of SP7-like effectors lacking their SP (ΔSP) fused with either their N-terminus (N-GFP) or C-terminus (C-GFP) to eGFP after transient expression in *N. benthamiana* epidermal leaf cells. **a** Shown are localization patterns in single plant nuclei. All but one effector target the plant nucleus. While RiSP7ΔSP exhibited a homogeneous nuclear distribution, RiSP2ΔSP, RiSP5ΔSP and the *Gigaspora* orthologs additionally localized at nuclear condensates. As exception, RiSP31ΔSP was mainly present in the surroundings of the nucleus (N-GFP) or detected only weakly in the nucleus (C-GFP). Scale bars represent 5 µm. **b** Representative overview pictures of SP7-like effector localizations (N-GFP) within the cellular context. As control, DsRed was co-expressed visualizing the plant nucleus and cytoplasm. All SP7 family effectors, with the exception of RiSP31ΔSP, are visible in plant nuclei (n, filled white arrowhead). In addition, RiSP7ΔSP, RiSP5ΔSP, RiSP31ΔSP and the *Gigaspora* proteins are present in the cytoplasm with varying intensities (cyt, blank arrowhead indicates exemplary region). No cytoplasmic signals can be detected for RiSP2ΔSP. ROI (Region of Interest, merged pictures) indicates starting points of transection lines used for fluorescence intensity measurements. Right: Fluorescence intensity blots along transection lines individually generated for GFP and DsRed channels. Nuclear areas and cytoplasmic signals at the cell periphery are boxed. Maximum fluorescence intensity = 255. **Note:** Only presence and changes in intensity at specific positions can be compared between two different fluorophores. Overall intensity levels can differ. Scale bar = 50 µm.

### SP7-like effectors highjack the mRNA processing pathway

In order to identify interaction partners of SP7-like effectors, we carried out yeast two hybrid (Y2H) assays using them as baits against libraries from several plants. Surprisingly, given the low overall sequence similarity among the fungal effectors, their Y2H interactome revealed a striking convergence on proteins related to mRNA processing, RNA binding and nuclear speckle categories, as revealed by gene onthology enrichment (Fig. 3a and Supplementary Table 3). These results are in line with the localization pattern observed in nuclear condensates and suggests that SP7-like effectors could be impacting on the fate of specific mRNAs. To obtain *in planta* evidence for this hypothesis, co-immunoprecipitation assays in *Nicotiana benthamiana* expressing GFP-tagged RiSP7 were carried out. The results not only validated several of the interaction proteins identified in the Y2H analysis, but also revealed further components of the mRNA processing from splicing to translation control (Fig. 3b, Supplementary Table 3).

**Figure 3.**
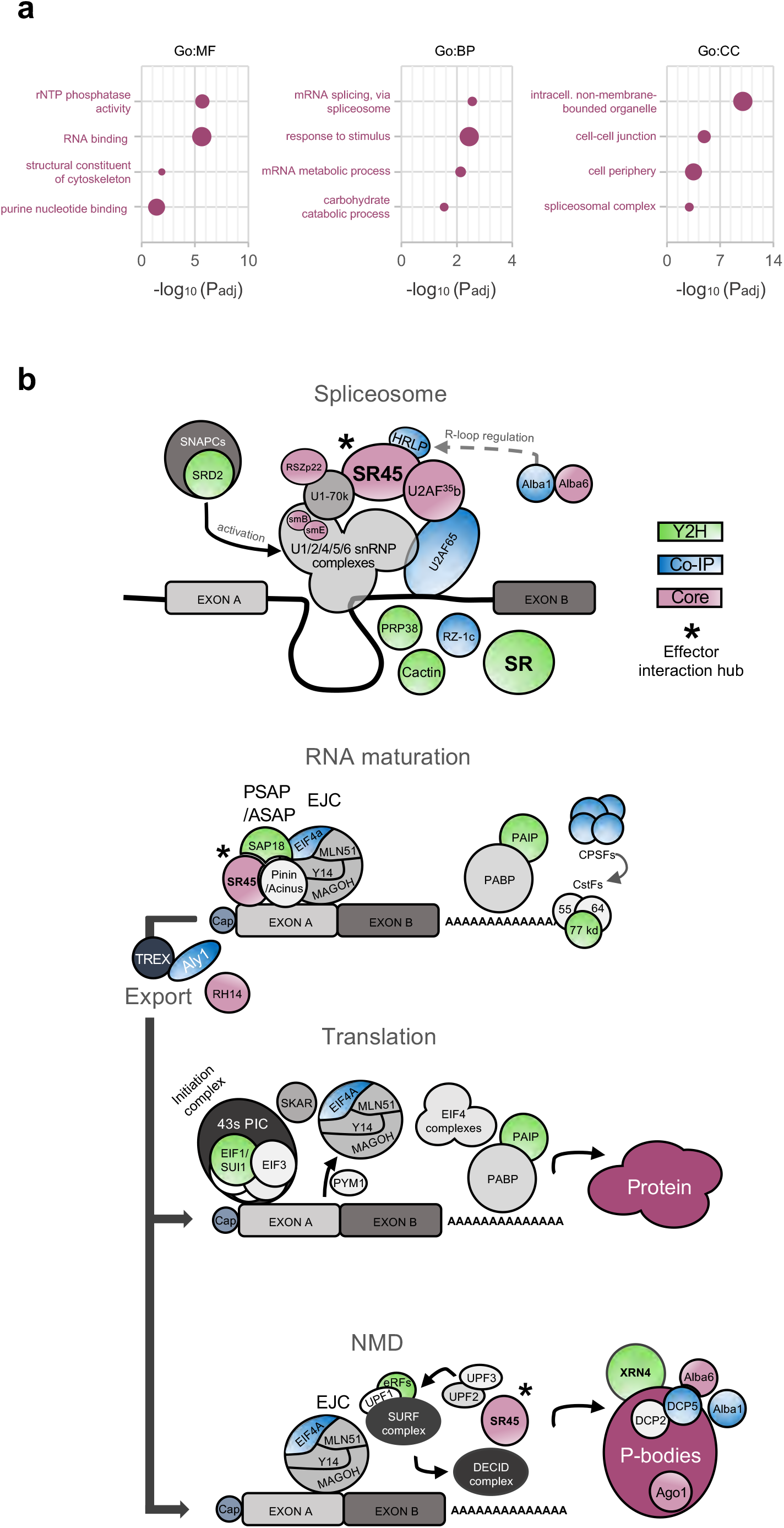
The interactome of SP7-like effectors is enriched for plant proteins involved in RNA metabolism. **a-b** By employing Y2H screens with SP7-like effectors as baits against *M. truncatula* and *A. thaliana* cDNA libraries as well as Co-IPs of RiSP7-GFP fusions (with and without SP) in *N. benthamiana*, a large set of putative plant interactors were identified. From those, 53 (core) effector target proteins either interacted with more than one effector or were repeatedly retrieved for RiSP7 in independent Co-IPs (detailed in Supplementary Table 3). **a** Go term enrichment analysis of the 53 core interactors using g:Profiler. Shown are the four most significant enriched functional gene groups in each category based on leading term filtering and p-value analysis (Supplementary Table 3). Significant enrichments of terms involved in RNA metabolism were found for each category. Circle sizes indicate number of interactors in each term. MF = Molecular function; BP = Biological process; CC = Cellular component. **b** Schematic illustration of a subset of identified interactors within the context of mRNA processing based on Arabidposis TAIR descriptions (Supplementary Table 3) and proposed functions (Woodward *et al*., 2017, Schlautmann and Gehring, 2020). With appearance in six different screens, SR45 (asterisk) was revealed as common effector interaction hub. Spliceosome: SnRNP complexes (activated by SNAPCs), SR proteins (including SR45/RNPS1) and other splicing factors assemble at splice sites. RNA maturation: After splicing, additional complexes are recruited at the exon junction including tetrameric EJC (exon junction complex), trimeric RNPS1/SR45 containing ASAP/PSPAP complexes and polyadenylation factors (PABP, PAIP, CPSFs). After nuclear export (TREX, Aly1), mRNAs are either translated or send for nonsense-mediated decay (NMD). Translation: EJC and peripheral factors dissociate (PYM1). Translation initiation complexes (43s PIC, SKAR) and ribosome association enable protein synthesis. NMD: RNPS1/SR45 and different NMD complexes (SURF, DECID) activate degradation of aberrant mRNAs in P-bodies. Green coloured = Y2H identified interactors. Blue coloured = Co-IP identified interactors (RiSP7). Red coloured = Core Interactors.

Interestingly, among SP7-like interactors, several serine-arginine (SR) rich proteins and their regulators were identified (Supplementary Table 3, Fig. 3b). Among those, the serine-arginine rich protein SR45 was consistently retrieved as interaction partner of all SP7-like proteins. SR proteins are often in multiprotein complexes regulating the different steps of mRNA life. For example, SC35, a central component that links SR proteins to co-transcriptional splicing^51^ was also identified as an interactor of RiSP2. This result suggests that SP7-like effectors could assemble with the spliceosome already during transcription. In support of this hypothesis, the largest subunit of the RNA polymerase II was immunoprecipitated with RiSP7 (Supplementary Table 3, Fig. 3b). Interestingly, the RNA binding protein HRLP, that together with SR45 inhibits the co-transcriptional splicing of intron I in the flowering repressor FLC, was also identified as interaction partner of RiSP7. HRLP leads to R-loop formation and transcription inhibition, thereby promoting flowering^52^.

Further support of the role of SP7-like effectors in mRNA splicing was obtained by the identification of two elements of the NineTeen Complex (NTC), MAC3a/Prp19 and MAC5a (Supplementary Table 3 and Fig. 3b). The NTC is a conserved eukaryotic protein complex that associates with the spliceosome and participates in splicing. In plants, it was shown to play a role in immunity as well as in developmental processes such as flowering^28,52^. Most interestingly, an interplay between RNA splicing and polyadenylation was also shown to play a role in immunity^28^. Here, we show that core components of the alternative polyadenylation pathway, such as CPSF30 or the cleavage stimulation factor CstF77 were also identified in the interactome of SP7-like effectors (Supplementary Table 3 and Fig. 3b). Altogether, these results suggest a picture where SP7-like effectors are recruited to nuclear condensates where they interact with components of the mRNA splicing machinery during transcription.

### SR45 is the central interaction hub of SP7-like effectors

Among all proteins identified in the interactome, only the SR protein SR45 was retrieved by all SP7-like effectors and was also immunoprecipitated by RiSP7 (Supplementary Table 3). Using targeted Y2H we could show that all SP7-like proteins interacted with the SR45 orthologues from three different plants, and with the two splice variants existing in *A. thaliana* (Fig. 4a). This is in agreement with the high degree of conservation of the plant SR45 proteins (Fig. 4b). MtSR45 from *M. truncatula* localizes in nuclear condensates (Fig. 4c) as it was previously shown for AtSR45^53^. Furthermore, both orthologues localize to the same condensates when co-expressed, while free eGFP does not (Fig. 4d). Bimolecular fluorescence complementation (BiFC) demonstrated that interactions between SP7-like effectors and MtSR45 also occurred in nuclear condensates (Fig. 4e). This was somewhat surprising, because while all SP7-like proteins showed a distinct nuclear localization when expressed alone, ranging from diffuse localization to accumulation at discrete nuclear bodies (Fig. 2a and Supplementary Fig. 4), co-expression with MtSR45, relocalized all SP7-like proteins to the same nuclear condensates occupied by MtSR45 (Fig. 4f). SR45 as well as RNPS1 contain two intrinsically disordered domains (Supplementary Fig. 5) which are thought to enable flexible interactions with different protein and RNA partners in a concentration-dependent manner^54,55^. Hence, effector re-localization might be triggered by an increased nucleating activity of MtSR45 when present at higher concentrations due to overexpression.

**Figure 4.**
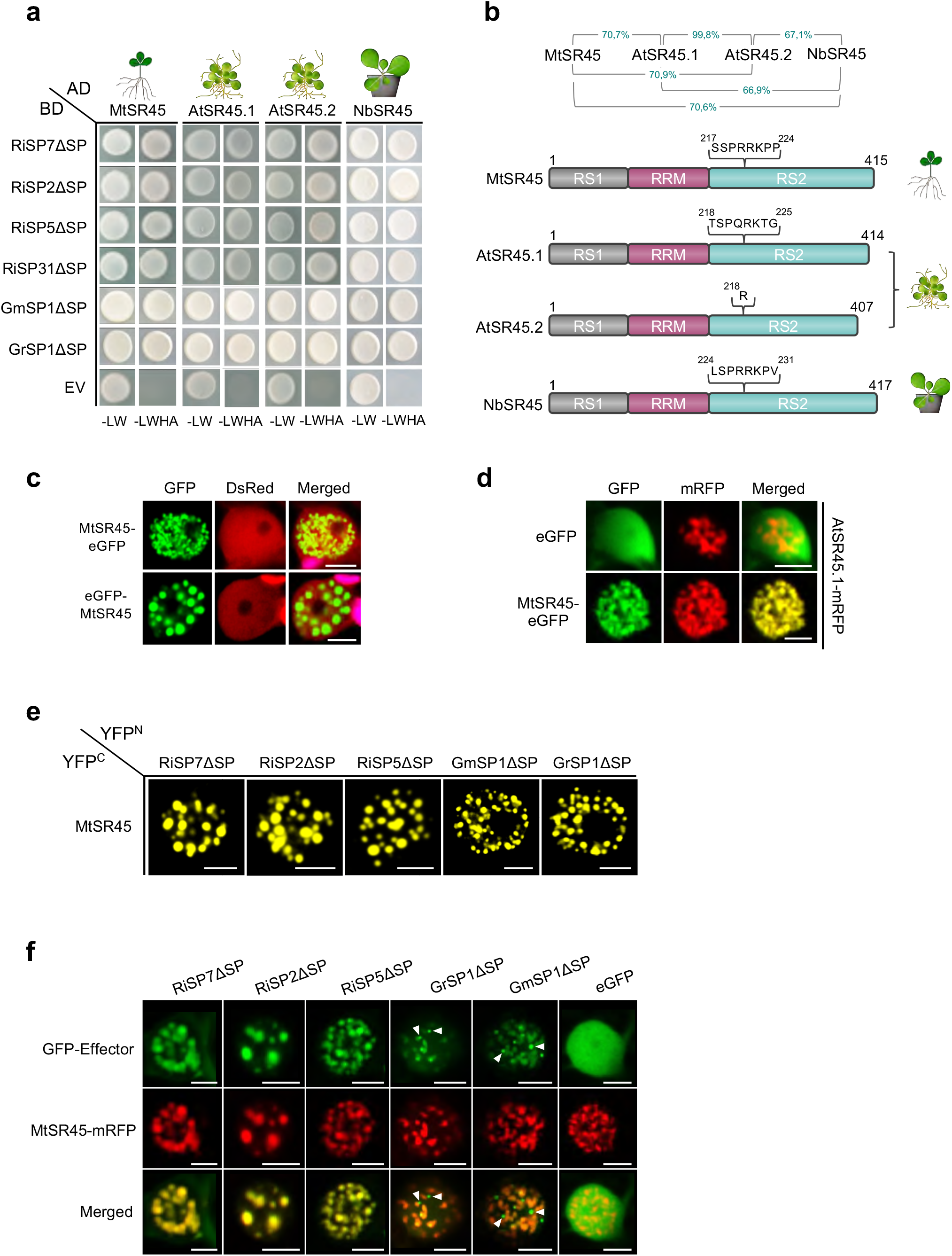
The SP7 effector family interacts with plant SR45 orthologs in nuclear condensates. **a** Direct yeast two hybrid interaction assays showing that all SP7-like effectors (Baits, BD) interact with plant SR45 orthologues from *M. truncatula*, *A. thaliana* and *N. benthamiana* (Preys, AD). EV = Empty vector. Positive interactions are indicated by colony growth on media lacking leucine, tryptophan, histidine, adenine (-LWHA). **b** Percentage protein identity (top) and protein structures of SR45 plant orthologs. SR45 proteins consist of a RS1, RRM and a RS2 domain. *AtSR45* codes for the two isoforms AtSR45.1 and AtSR45.2 (Zhang and Mount, 2009). **c-f** Shown are localization patterns in single plant nuclei after transient expression of fluorescent tagged proteins in *N. benthamiana* epidermal cells. Scale bars represent 5 µm. **c** Subcelluar localization of MtSR45 (N- or C-terminal eGFP fusion) in different shaped nuclear speckles. As control, free DsRed was co-expressed. Chloroplast autofluorescences are pseudo-coloured in magenta (merged pictures). **d** Co-localization of MtSR45 C-terminally fused to eGFP and AtSR45.1 C-terminally fused to mRFP. MtSR45 and AtSR45.1 co-localize in nuclear speckles. As control, free expressed eGFP does not accumulate in speckles. **e** Bimolecular fluorescence complementation assays to confirm yeast two hybrid results from (a). Reconstituted YFP fluorescence was observed in nuclear condensates for plants co-expressing MtSR45 (fused to C-terminal half of YFP, YFP^C^) with all tested SP7-like effectors (fused to N-terminal half of YFP, YFP^N^). **f** Co-localization of SP7-like effectors N-terminally fused to eGFP and MtSR45 fused C-terminally to mRFP. In contrast to the eGFP control, all tested SP7-like effectors re-localized to nuclear condensates occupied by MtSR45. For *Gigaspora* effectors, exclusive nuclear bodies were observed in GFP channels (white arrowheads). **a-f** All experiments were carried out with SP7-like effectors lacking their signal peptides (ΔSP).

To map the SR45 domain required for the interaction with SP7-like effectors, we next analyzed the ability of truncated versions of MtSR45 to interact with RiSP7 in a Y2H screen. The results showed that the carboxy terminal RS2 domain of MtSR45 is necessary and sufficient to mediate the interaction with RiSP7, while the RNA binding and the RS1 domain are dispensable (Fig. 5a). This result was confirmed for all SP7-like proteins and corroborated by BiFC assays (Fig. 5b, c and Supplementary Fig. 6a, b). This finding is interesting, because the RS2 domain had been shown to be necessary and sufficient to drive localization of AtSR45 to nuclear condensates^56^. We showed that this is also the case for the *M. truncatula* orthologue (Supplementary Fig. 6c). Furthermore, SP7-like effectors only fully co-localize with MtSR45 if the RS2 domain is present and the protein forms nuclear condensates (Fig. 5d and Supplementary Fig. 6d).

**Figure 5.**
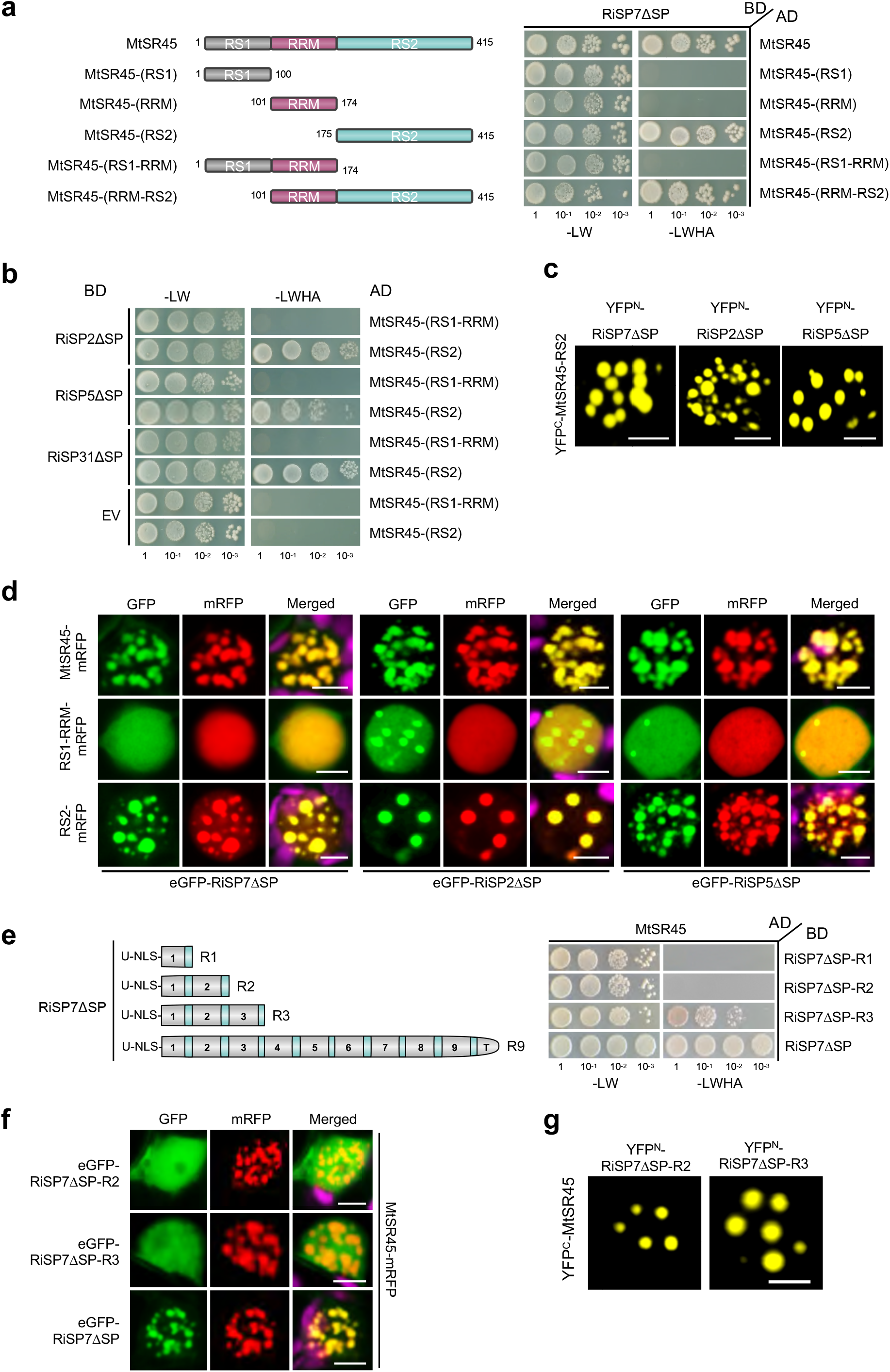
The RS2 domain of MtSR45 is required and sufficient for effector interaction and number of repeats determine interaction strength. **a-f** All Y2H spotting assays were performed using different yeast dilution series (1-10^-3^) with SP7-like effector constructs as prey (AD) and MtSR45 constructs as bait (BD). Positive interactions are indicated by colony growth on media lacking leucine, tryptophan, histidine, adenine (-LWHA). EV = Empty vector. For localizations and Bimolecular fluorescence complementation (BiFC) assays, single plant nuclei are shown after transient expression of fluorescent tagged proteins in *N. benthamiana* epidermal cells. Scale bars represent 5 µm. Experiments were performed with SP7-like effectors lacking their signal peptides (ΔSP). **d, f** Chloroplast autofluorescences are pseudo-coloured in magenta (merged pictures). **a** Interaction assay of RiSP7 with different MtSR45 domains. Left side: Overview of generated MtSR45 truncation constructs. Right side: Y2H using RiSP7ΔSP against MtSR45 and truncations. Only proteins containing the RS2 domain interacted with RiSP7ΔSP. **b** Y2H as described in (a) with additional SP7-like effectors. **c** Confirmation of RS2 domain as effector target site using BiFC (RS2 domain fused to C-terminal half of YFP, YFP^C^, SP7-like effectors fused to N-terminal half of YFP, YFP^N^). **d** Co-localization of RiSP7ΔSP, RiSP2ΔSP or RiSP5ΔSP (N-terminal GFP fusion) with C-terminal mRFP fusions of MtSR45, RS1-RRM or RS2 domain alone. Only expression of RS2 domain containing proteins led to co-localizations into nuclear condensates. **e** Interaction assay of truncated RiSP7 versions. Left side: Overview of generated RiSP7ΔSP truncations (Full length = R9, 1-3 repeats = R1-R3). Right side: Y2H using RiSP7ΔSP truncations against MtSR45. **f** Co-localization of RiSP7ΔSP truncations (N-terminal GFP fusion) with MtSR45-mRFP. Minimum repeat number for effector re-localization into MtSR45 nuclear condensates was three. **g** BiFC of RiSP7ΔSP-R2 truncation (fused to N-terminal half of YFP, YFP^N^) and MtSR45 (fused to C-terminal half of YFP, YFP^C^) showed interaction in nuclear condensates.

As shown above, SP7-like protein effectors share only a low similarity regarding their overall sequence identity, but all follow the same modular protein structure. All the repeat subunits, except the truncated last repeat, end in a small stretch of highly conserved amino acids (Fig. 1a, Supplementary Fig. 1a). Given that the number of repeats in each of the SP7-like proteins is variable, we next asked the question of how many protein repeats were required for the interaction with SR45. To answer that, truncated versions of the RiSP7 protein containing one, two or three repeats were tested for their ability to bind MtSR45. When the interaction was analyzed by Y2H, a minimum of three repeats were required, but this interaction was not as strong as with the full-length RiSP7 protein (Fig. 5e). Experiments in *Nicotiana* showed that full co-localization with MtSR45 in nuclear condensates was only observed with the full-length protein, while three repeats were sufficient to show a partial co-localization, supporting the Y2H results (Fig. 5f). Surprisingly, a two repeats construct was sufficient to allow BiFC (Fig. 5g and Supplementary Fig. 7a), suggesting that the interaction between RiSP7 and MtSR45 could be mediated by small linear motifs between each RiSP7 repeat and the RS2 domain of MtSR45, and that the strength of the interaction increases with the number of repeats (Supplementary Fig. 7b).

### The splicing proteins U2AF^35^b and U1-70K are also core interactors of SP7-like effectors

Arabidopsis SR45 (Fig. 6a) was originally discovered as an interacting protein of the splicing factor U1-70K and later also shown to interact with the splicing factor U2AF^35^b^40,57^. U2AF^35^b was already retrieved as interacting partner of RiSP7 when the effector was first identified^12^ and now in our interactome here. Both splicing factors were shown not only to interact with AtSR45 but also to co-localize in the nucleus^40,56^. MtSR45 did also interact with both splicing factors in yeast and in *Nicotiana* and localized to the same nuclear condensates (Supplementary Fig. 8). We then tested whether SP7-like effectors were able to directly interact with both splicing factors. Indeed, this was the case, and all effectors interacted with MtU1-70K and MtU2AF^35^b, both in Y2H as well as in *Nicotiana* assays (Fig. 6b and c). Interestingly, the reconstituted YFP signal in BiFC assays between RiSP7-MtU1-70K and the RiSP7-MtU2AF^35^b interactions, was observed as a nucleoplasmic signal which was often accompanied by a low number (usually one to four) of nuclear foci (Fig. 6c). In contrast, the interactions between all other SP7-like effectors was observed mainly in nuclear condensates (Fig. 6c). This suggests, that in contrast to MtSR45, the splicing proteins MtU1-70K and MtU2AF^35^b have a weaker effect as nucleating factors.

**Figure 6:**
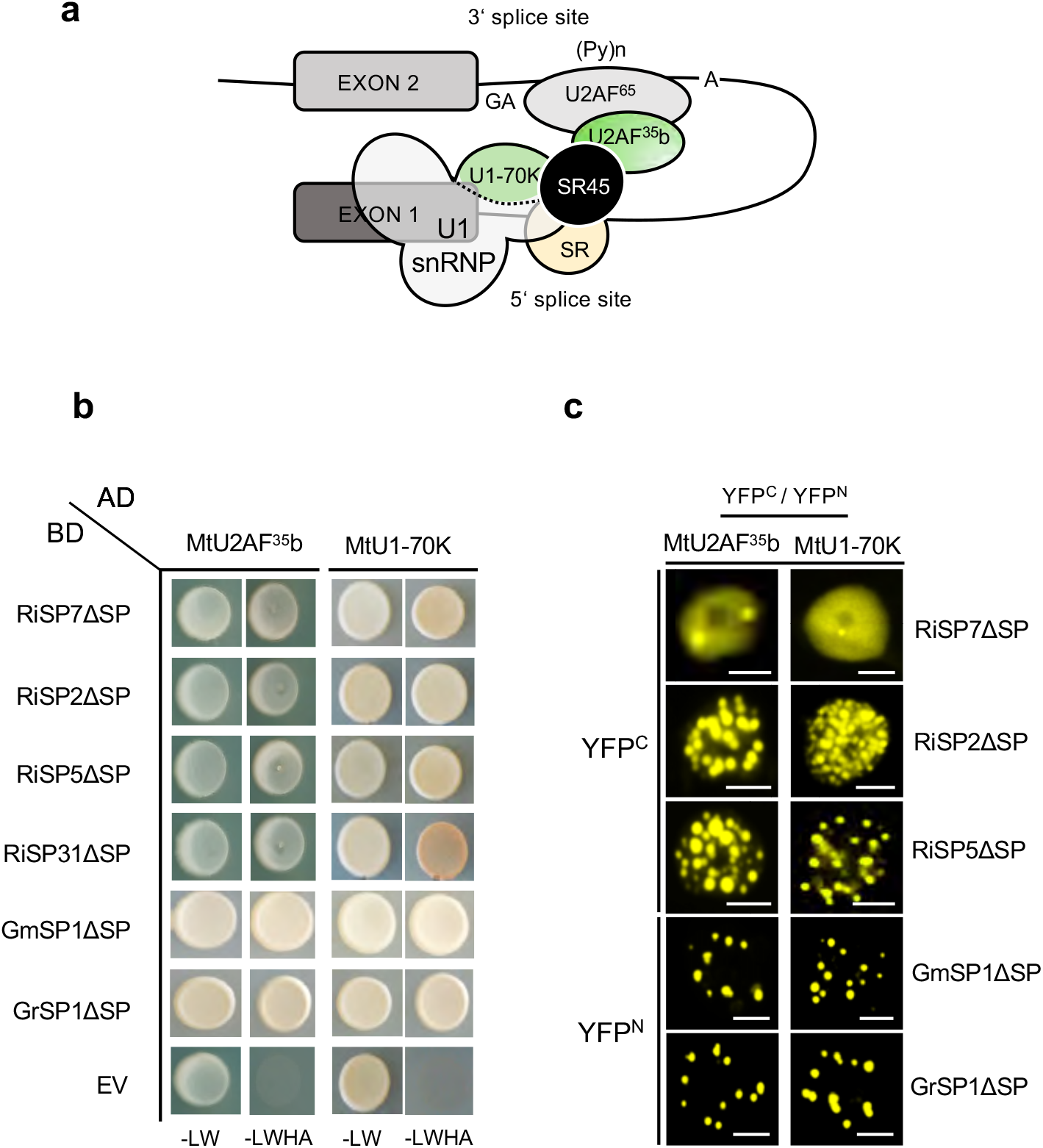
SP7-like effectors interact with SR45 associated spliceosomal components U1-70K and U2AF^35^b. **a** Illustration of known interaction network between *Arabidopsis* SR45 and spliceosomal proteins at early stages of pre-mRNA splicing. AtSR45 directly interacts with U1-70k (part of U1 snRNP) and U2AF^35^b (forms heterodimer with U2AF^65^) at splice sites (Golovkin and Reddy, 1999; Ali *et al*., 2008; Day *et al*., 2012). This interaction network may serve as bridging platform to facilitate splice-site pairing across introns and to recruit components for spliceosome complex formation (Day *et al*., 2012). The dual interaction of SR45 with U1-70k and U2AF^35^b is conserved in *Medicago* (see Supplementary Fig. 9). GA = 3’ splice site, (Py)n = Polypyrimidine tract, A = branchpoint adenosine, SR = additional SR proteins. **b** Direct Y2H assays showing that all SP7-like effectors (BD) interacted with *Medicago* U2AF^35^b and U1-70k spliceosome components (as Preys, AD). Positive interactions are indicated by colony growth on media lacking leucine, tryptophan, histidine, adenine (-LWHA). EV = Empty vector. **c** Validation of interactions seen in (a) by BiFC assays in *N. benthamiana* leaf epidermal cells. Shown are magnified pictures of single plant nuclei. Reconstituted YFP fluorescence was detected in nuclear condensates for plants co-expressing either MtU2AF^35^b or MtU1-70k with all tested SP7-like effectors. For RiSP7ΔSP only a few nuclear condensates could be observed. Instead, YFP signals were mainly distributed evenly throughout the nucleoplasm. For MtU2AF^35^b and MtU1-70k N-terminal fusions of the N-terminal half of YFP (YFP^N^) were used for co-expression with N-terminal fusions of RiSP7ΔSP, RiSP2ΔSP and RiSP5ΔSP (fused to C-terminal half of YFP, YFP^C^), while for experiments with *Gigaspora* members the YFP fusion halfs (YFP^N^, YFP^C^) were interchanged. Scale bars represent 5 µm. All experiments were performed with SP7-like effectors lacking their signal peptides (ΔSP). Alteration of CDS (Stop codon conserved)

### SP7 ectopic expression induces alternative splicing in potato

Given that MtSR45 and MtU1-70K and MtU2AF^35^b are central component of the interactome of SP7-like effectors, it seems plausible to think that the function of these effectors might be to alter the fate of specific target mRNAs by inducing alternative splicing. To challenge this hypothesis, we constitutively expressed RiSP7 in potato plants and analyzed its effect on alternative splicing in roots. Our results suggest that ectopic expression of the effector protein RiSP7 leads to alternative splicing events in a small subset of genes (96) (Supplementary Table 4) within the whole landscape of AS genes (7161) from potato (Phytozome v13). From the 96 DAS (differential alternative splicing) genes identified in response to the ectopic expression of SP7, eight were also differentially expressed with seven of them repressed (Supplementary Table 4), suggesting that their AS forms might be also subjected to non-sense mediated decay. However, the majority of DAS genes did not show an overall change in expression. Among the alternative splice events observed in response to ectopic expression of RiSP7, most of them corresponded to intron retention (44%) followed by changes in alternative 3’ or 5’ splicing site (Supplementary Table 4).

In order to get insights in the involvement of SR45 in the SP7-mediated alternative splicing, we compared our data set with that of two other studies in *A. thaliana*^26,58^, where SR45 mRNA targets were identified by RNA immunoprecipitation (Fig. 7a). We could identify 87 putative Arabidopsis orthologues of the SP7-mediated potato DAS genes (Supplementary Table 4). From those, 48 were also identified as AtSR45 mRNA targets (Fig. 7a and 7b). To confirm the alternative splicing results, we randomly selected 11 of those genes showing different types of AS events and analyzed the expression of the different variants using qRT-PCR with specific primers (Fig. 7c). From those genes, most of them (6) had splicing events that altered the UTR regions or produced premature stop codons (4), and only one gave rise to an alteration of the coding sequence without affecting the stop codon. We confirmed that 9 of those genes underwent alternative splicing in response to RiSP7. Towards evaluating whether these alternative splicing events were also induced by other SP7-like effectors, transgenic potato plants expressing RiSP5 were generated. Strikingly, 6 from the 11 analyzed genes in RiSP7 transgenic potato were also alternatively spliced in response to the ectopic expression of the RiSP5 fungal effector (Supplementary Fig. 9), suggesting a conserved mechanism for this effector family to modify alternative splicing in plants.

**Figure 7.**
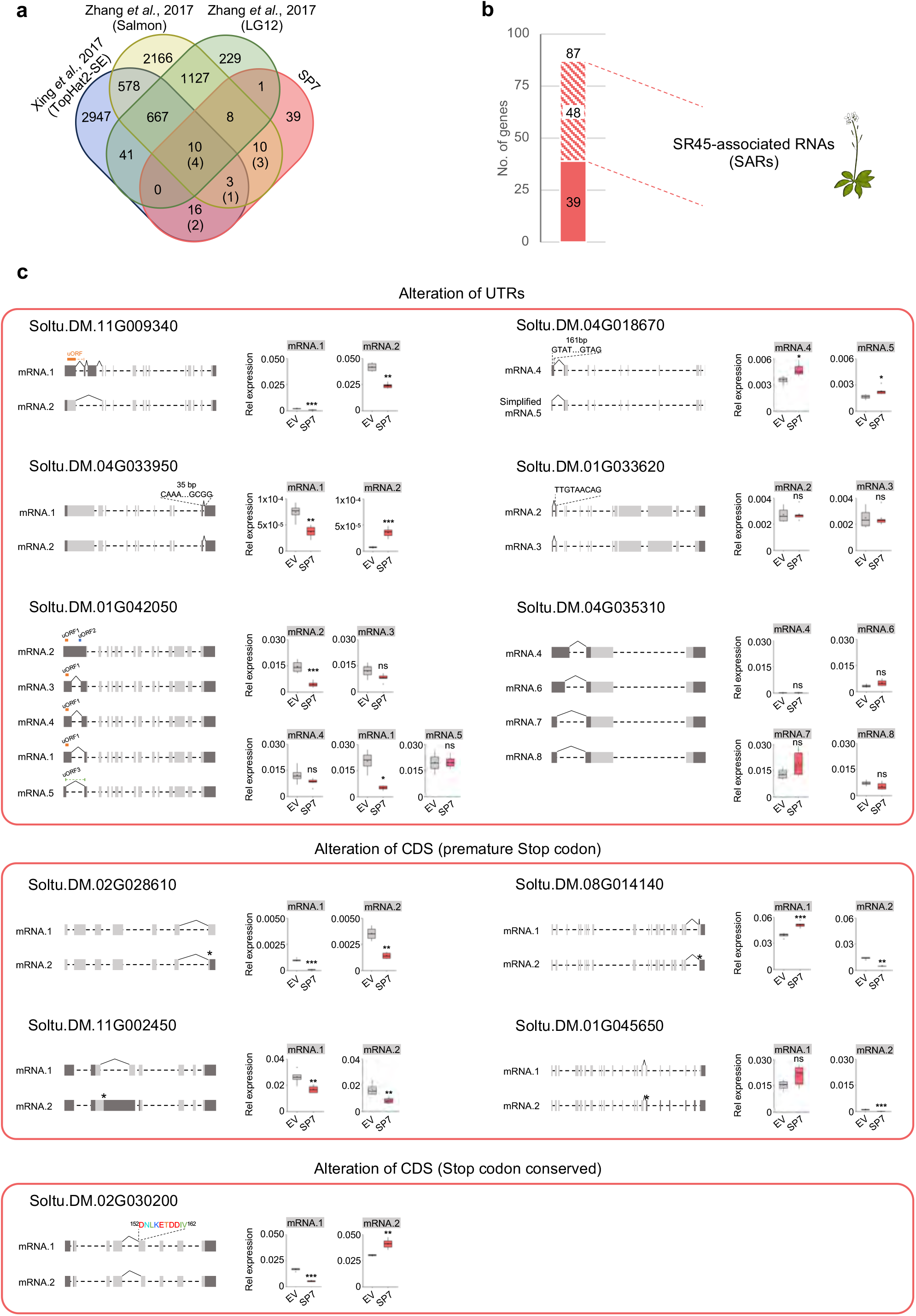
The extopic expression of SP7 alters the plant’s alternative splicing landscape. **a** Venn diagram showing the overlaps of the SR45 associated RNAs (SARs) from *Arabidopsis thaliana* found in the works of Xing *et al.* 2015 (TopHat2-SE pipeline) and Zhang *et al.* 2017 (Salmon and LG12 pipelines) with the *Arabidopsis* orthologs of the potato genes whose alternative splicing was regulated upon overexpression of RiSP7. In brackets is shown the amount of genes selected for verification of the splicing analysis. **b** Stacked column chart representing the *Arabidopsis* orthologs of the potato SP7-regulated DAS genes from (a) and how many of those are found as SARs. **c** Splicing models and quantification of the relative expression of each splicing isoform through qRT-PCR of a random selection of SP7-regulated DAS genes that also are identified as SARs. In the splicing models the dashed lines represent introns and the boxes exons. Boxes in dark grey coorespond to untranslated regions (UTRs) and those in light grey to protein coding sequence (CDS). Expression values relative to *StActin* are shown as boxplots. Statistical significance was assessed by performing a Student T test or a Mann Whitney U test after checking the normal distribution of the data and homocedasticity as explained in Materials and Methods. Significance levels: n.s. (non-significant) p>0.05, * p<0.05, ** p<0.01, *** p<0.001.

## Discussion

Plants are constantly challenged by interacting microbes and adapt their physiology modulating their gene expression accordingly. In recent years it has become evident that part of the plant physiological response to microbes also involves alternative splicing as a posttranscriptional mechanism to allow fast adaptation to the environment^24^. Although the mechanisms are starting to be elucidated, it has been shown that in several pathogenic interactions, these changes in AS can be directly caused by the microbe through the delivery of effector proteins, mainly with the aim to modulate immunity. Here we show that symbiotic arbuscular mycorrhizal fungi have evolved an effector family, the SP7-like family, able to interact with the plant mRNA processing machinery and induce AS of a small subset of genes. The funder member of this effector family, SP7, was identified in *R. irregularis* using a genetic screen for secreted proteins that contained an NLS in their sequence^12^. There, we could already identify other paralogues having a similar protein structure (hydrophylic tandem repeats) but only SP7 was functionally analyzed and shown to localize in the nucleus. The release of the first *R. irregularis* genome revealed several other putative paralogues^7,8^, but only now, with the publication of a largely improved sequence and annotation^9,10^, the full SP7-family could be disclosed. Our phylogenetic analysis shows that the SP7-like family seems to be unique to the Glomeromycotina and widespread among different orders. However, the fact that we could not find evidence of this family in the orders Paraglomerales or Archaeosporales suggests that the ancestor of this gene family might have not been present in the earliest AM fungi^59^.

We had previously shown that RiSP7 localizes at the nucleus, and that its ectopic expression in roots is able to promote biotrophy and counteract defense responses^12^. The identification of novel members in the SP7-like family lacking a predicted NLS challenged our hypothesis that these proteins would be nuclear effectors modulating immunity. However, localization assays *in planta* of different members of the family revealed that they all targeted the nucleus. This might be explained by the presence of the basic amino acids KR in the conserved signature present in each repeat (Fig.1) or other non-conventional NLS^17^. The amino acid analysis of SP7-like effectors revealed that their composition is biased towards the presence of five specific amino acids (Asp, Tyr, Lys, Ser and Gly) that represents between 75 and 80% of their sequence (Supplementary Figure 10). This is in agreement with their nature as proteins containing large intrinsically disordered regions (IDRs) which are characterized as having low sequence complexity and being enriched in a limited number of amino acids such as those in SP7-like proteins^49^. Proteins with large IDRs usually lack a defined 3D structure but are often organized in repeat elements that allows multivalent intermolecular interactions. They are common components of biomolecular condensates, particularly those in which RNA accumulates such as nuclear condensates, P bodies or stress granula^49^. This localization is due to the low amino acid diversity that promotes the appearance of blocks of positively and negatively charged amino acids that are critical for targeting to RNA granules. This is consistent with the observed localization of SP7-like proteins in nuclear condensates and also in P bodies.

Biomolecular condensates can mediate specific reactions or increase the dwell time by bringing molecules together or serve the sequestration of specific proteins in plants^60^, where they are increasingly recognized as playing a role in immunity^61^. Thus, the localization of SP7-like proteins and their interactome supports a role for these effectors in mRNA processing. We hypothesize that they are able to do so by bringing together specific mRNAs and RNA processing proteins. This is also consistent with the ability of proteins having IDRs to drive multivalency, the ability to interact with multiple partners through several interaction sites, forming a highly connected network through noncovalent crosslinks with other proteins^49,50^. These interactions might be weak individually what allows for liquid-liquid phase transitions^49,50^ as observed when analyzing the strength of the interaction between single or multiple repeats of RiSP7 and MtSR45.

The fate mRNAs is strongly determined by the set of RNA binding proteins that accompany the transcript from early stages of mRNA biogenesis until their translation, from which SR proteins play a major role^35,36^. Our interactome results show that SP7-like effectors appear associated with many components of this pathway, and most prominently with the unconventional SR protein SR45 and the splicing factors U2AF35b and U1-70K. Taken together, this suggests that impacting on alternative splicing might be a key function of this effector family. However, other identified components suggest that these effectors might not be involved only in early stages of mRNA processing but also accompany the mRNAs and SR45 through their journey to translation. This is not totally surprising since SR proteins have sometimes been found to remain associated to specific mRNAs and participate in their export out of the nucleus and in their translation^35^. In fact, most SR proteins shuttle between the nucleus and the cytoplasm, with some of them being export adaptors for specific mRNAs out of the nucleus, such as SRSF3 and SRSF7^62^. RSZ22, the orthologue of SRSF7, and involved in immunity^63^, was retrieved as interactor of two SP7-like proteins (Supplementary Table 3, Fig. 3b). Furthermore, other proteins controlling the nuclear-cytoplasmic shuttling such as the chaperons HSP90, 70, 40 and 60^64^, the specific SR protein kinases such as SRPK1^64^, the helicase DRH1 required during defense responses^65^, or ALY1, a component of the TREX complex^66^ were also identified in the SP7-like interactome. And even more, several ALBA proteins known to bind specific mRNAs and control their translation by phase separation between stress granula and P bodies^67^ were identified in the interactome of SP7-like effector. Altogether, we postulate that SP7-like effectors are delivered to the nucleus to alter the fate of specific mRNAs by interacting with the plant mRNA machinery to alter their splicing and/or their translation.

In this context, the main interactor of SP7-like effectors, the SR45 protein is a crucial component. SR45 is the orthologue of the well-studied human RNPS1 protein that plays a dual role in splicing and mRNA quality control^41,42^. The fact that all SP7-like proteins investigated interacted directly also with the core splicing proteins U1-70K and U2AF35 suggest that their function begins at the onset of splicing possibly already during transcription. RNA immunoprecipitation assays with SR45 in *Arabidopsis* have shown that more than 4000 genes are associated with this protein, including intron-less genes^58^. Given that SR45 has been involved in many developmental, stress and immunity related processes^26,37–39,43,68^, explaining the pleiotropic phenotypes of the *sr45-1* mutant, it is clear that target specificity must be given by other RNA binding regulators.

Alternative splicing is increasingly recognized to play a key role in controlling innate immunity in plants, as many mutants in splicing components show an immunity related phenotype^23^. For example, CPR5, an *Arabidopsis* RNA binding protein from the serine-arginine rich family that localizes in nuclear speckles, has been recently shown to modulate plant immunity. It does so by interacting at those speckles with two key regulators of RNA processing, the NineTeen Complex and the cleavage and polyadenylation specificity factor, to control the AS of more than 500 transcripts^28^. Interestingly, both of those components were found as interactors of SP7 (Supplementary Table 3). It is thus not surprising that AS has been shown to be a target for several effector proteins in order to subvert plant immunity^31,32,51^. Because most of the splicing events observed were not paralleled to changes in transcript levels of the genes affected, we assume that the production of proteins with different sequence or domain arrangements might be the main function of SP7-like proteins. This might serve different purposes such as producing different subcellular localization, changes in protein stability, alteration of translation speed or even producing dominant-negative versions of the target proteins.

There are several examples of microbial effectors that can target components of the host mRNA processing, but only a few of them have been shown to target host alternative splicing. One of them is HopU1 from the bacterial pathogen *Pseudomonas syringae*. This effector had been identified as an ADP-ribosylation factor of RNA binding proteins, GRP7 and GRP8, interfering with immunity^33,69^. GRP7 can alter the choice of alternative 5‘ splice sites of specific mRNAs^70^. Further studies revealed that in the case of GRP7, HopU1 impaired immunity by preventing the interaction of GRP7 with the mRNA of immune receptors such as FLS2 and EFR and decreased their protein abundance. However, it was not clear if this was achieved by an alternative splicing mechanism^71^. In contrast, several other studies have directly involved effectors in the modulating AS in their plant hosts, including cyst and root-knot nematodes^34^. MiEFF18 is an effector from *Meloidogyne incognita* that interact with the spliceosomal protein SmD1 and modifies AS impacting on giant cell formation^34^. Interestingly, the orthologue of SmD1 was also immunoprecipitated by SP7 (Supplementary Table 3). Also, two seminal studies from Huang and collaborators^31,32^ revealed that alternative splicing by effectors is an important mechanism to reprogram plant immunity in the oomycete pathogen *Phytophthora* (from 87 RXLR effectors, 9 regulate splicing). One of them, PsAvr3c, interacts with SKRP-like proteins and alter alternative RNA splicing of more than 400 genes. SKRP proteins turned out to be negative regulators of plant immunity, but most interestingly immunoprecipitation assays showed an interaction with SR45 and with U1-70K and U2AF35 suggesting that they are part of the spliceosome. They could show that both PsAvr3c and SKRP ectopic expressions induced alternative splicing and that SKRP targets exon 3’ end of unspliced RNA^32 68^. This reveals a similar picture to the function of SP7-like effectors and their interactors, although the amount of alternative spliced genes in response to ectopic expression of the mycorrhizal effector is much lower (20 times). This is also not surprising, because in contrast to the large amount of alternative spliced plant genes observed in response to Phytophthora infection, more than 5000 genes in tomato^31^, the number of alternative spliced genes during arbuscular mycorrhiza colonization was shown to be very low, with only 500 events in pea roots^72^. Although we believe that certainly the number of AS during mycorrhizal symbiosis is lower than in the case of *Phytophthora*, we think that this number is somewhat falsified due to the dilution effect. We expect most AM fungal effectors *in planta* to be expressed and secreted at the major symbiotic interface, the arbuscule^15^. Thus, the analysis of whole roots might obscure the real number of AS transcripts. Furthermore, the root colonization by AM fungi is not a synchronous process, and therefore, analyzing whole roots might also not show the dynamic variation in AS during the colonization process.

Our results here demonstrate that the family of SP7-like effectors from AM fungi localize to nuclear condensates and interact with components of the plant mRNA processing pathway at those loci. We propose that they act to modify the AS program of their host plants and/or the fate of specific mRNAs. Although among the potato AS genes in response to RiSP7 there are some candidates that could be involved in reprogramming the immune system of the plant and prevent defense reactions, we need further work to analyze and confirm this hypothesis. Furthermore, it is possible that other plant developmental or nutritional processes are targeted by this effector family that could indirectly impact on the immune system of the plant. Understanding the mechanisms of communication between mutualistic symbionts and their crop plants will help to identify targets that can be engineered for mechanisms of resilience and improved nutrient acquisition.

## Methods

### Bioinformatic online resources used in this study

Signal peptide prediction for SP7-like effector proteins was carried out using SignalP5.0, (https://services.healthtech.dtu.dk/services/SignalP-5.0/)^73^. The presence of possible NLS in SP7-like effectors was predicted using the traditional PSORTII prediction of WoLFPSORT (https://wolfpsort.hgc.jp/)^74^ with plant as the selected organism type. Disordered regions in SP7-like effectors and MtSR45 were predicted using PrDOS, (http://prdos.hgc.jp/cgi-bin/top.cgi)^75^ and IUPred3 (https://iupred3.elte.hu/plot) ^76^ with default parameters. Alignments and comparative percentage identity analysis were performed using Clustal Omega (http://www.ebi.ac.uk/Tools/msa/clustalo/)^77^. For analysis of consensus motifs, highly conserved regions of always the second repeat from all 21 identified full length SP7-like effectors (Supplementary Table 2, Supplementary Figure 1a) were manually aligned and visualized with ESPpript 3 (http://espript.ibcp.fr/ESPript/ESPript/)^78^. Aligned repeat regions were further used in WebLogo (http://weblogo.berkeley.edu) ^79^, to create the consensus logo with the effector family characteristic Y[KR]Rx[AP]x[AP] motif. Protein, gene and *R. irregularis* chromosome models were scaled using IBS^80^ and redrawn according to these scales. The evolutionary relationship of SP7-like effectors was inferred by using the Maximum Likelihood method and JTT matrix-based model using MEGA11^81^ with 1000 bootstrap replicates. For that purpose, the protein sequences of all 21 identified full length SP7-like effector members were first aligned using ClustalW in MEGA11. Initial trees for the heuristic search were obtained automatically by applying Neighbor-Joining and BioNJ algorithms to a matrix of pairwise distances estimated using the JTT model, and then selecting the topology with superior log likelihood value. The tree with the highest log likelihood is shown (Fig. 1c) and visualized using iTOL (https://itol.embl.de/)^82^. Bootstrap values are indicated by node circles. Branch lengths are ingnored. To identify SP7-like genes in Mucoromycota species, genome data for Glomeromycotina, 19 different Mucoromycotina and Mortierellamycotina species as well as *vermifera* were BLAST searched using BlasterQt^83^ to create a local BLASTp interface. Protein sequences and motifs of known ^7,8,12^ SP7-like effectors were used as query. BLASTp hits and respective gene models thereof were further analyzed for the existence of predicted signal peptides, repeats, conserved motifs and introns. Identified SP7-like effector protein and gene sequences are summarized in Supplementary Table 2. To assess whether SP7-like effector transcripts are present in AM species and different fungi outside the Glomeromycotina, effector coding sequences were used as query to perform BlastN analysis in SRA (Sequence Reads Archive) databases. For that purpose, publicly available RNA based experiments (Filter: RNA) from organisms belonging to Mucoromycotina, Blastocladiomycota, Zoopagomycota, Mortierellomycotina and Glomeromycotina were separately selected and used for Blast analysis with the parameter “more dissimilar sequences”. For Chytridiomycota (*Batrachochytrium dendrobatidis*), Ascomycota (*Blumeria graminis*, *Laccaria bicolor*), Basidiomycota (*Uromyces spp*., *Serendipita vermifera*) and *R. irregularis* exemplary experiments were selected. All obtained reads are listed and compared in Supplementary Table 1. In all cases, the criteria used to assign reads as significant matches were a) they would show at least 75% identity within the alignment; b) an alignment length should be over 94 bp; and c) at least one of these reads spans the SP7-like family consensus motif after translation. Go term enrichment analyses of identified plant interaction partners were performed using g:GOSt functional profiling at the g:Profiler web service^84^.

### Biological material and transformation of plants

Leaf disk transformation of potato (*Solanum tuberosum,* cv. Desiree) was carried out using *Agrobacterium tumefaciens* (AGL-1 strain) with overexpression constructs for RiSP7, RiSP5 or an empty vector (EV). Transgenic plants were grown in a sand-gravel mixture (4:1) at 21°C *N. benthamiana* plants were grown in soil at 28°C. All plants had a light cycle of 16 h day/8 h night and were grown in a GroBanks Birghtboy-XXL.2 (CLF Plant Climatics, Wertingen, Germany).

*Rhizophagus irregularis* DAOM 197198 was cultivated in monaxenic culture as described before^85^. *Gigaspora margarita* (BEG34) spore inoculum was obtained from Dr. J. Palenzuela (EEZ, CSIC, Spain) and *Gigaspora rosea* (DAOM194757) spore inocula were donated by Dr. C. Roux (CNRS, Toulouse, France) and from Dr. S. Roy (Agronutrition, France).

### Genomic DNA isolation from fungal spores

Spores of *R. irregularis* DAOM 197198 (approximately 1000), *G. margarita* (BEG 34) or *G. rosea* (DAOM194757) (for both approximately 300 spores) were surface sterilised for 30 min (4% Chloramine T) and incubated in a 100 µg ml^−1^ gentamycin and 200 µg ml^−1^ streptomycin antibiotic solution. After washing with sterile _d_H_2_O spores were crushed in 0.1% sarkosyl in TE-buffer solution using a pestle for eppendorf tubes. Pronase (0.25 mg ml^−1^) and 1% SDS were added to the spore solution and incubated at 65 °C for 1 h. Samples were homogenized every 15 min. Subsequently 1M NaCl was added and solutions were incubated on ice for 30 min. Genomic DNA was then extracted and precipitated using phenol/chloroform and standard protocols.

### RNA isolation from plant and fungal material

For mRNA extraction frozen plant or fungal material was homogenised with 5 mm metal beads in a cell mill MM200 Retsch. The frozen material was homogenised in three cycles of 1 min at a frequency of 25/s, and refrozen in liquid nitrogen between each cycle. Plant and fungal mRNA was extracted using the innuPREP Plant RNA kit (Analytik Jena, Jena, Germany) or the TRIzol method (ThermoFischer Scientific, Waltham, MA, USA) according to manufacturer instructions.

### Molecular cloning and vector plasmid construction

The genomic or cDNA sequences of SP7-like effectors *RiSP7*, *RiSP2*, *RiSP5*, *RiSP31*, *GmSP1*, *GrSP1* were amplified and cloned in pCR2.1/TOPO, pCR8/GW/TOPO or pENTR/D-TOPO (ThermoFischer Scientific, Waltham, MA, USA) with the primers listed in Supplementary Table 5. *MtSR45* (*Medtr2g069490*), *MtU1-70K* (*Medtr8g077840*), *MtU2AF35b* (*Medtr1g035130*), *NbSR45* (*Nb101Scf00283g0008*), *AtSR45* (*At1g16610*), *AtDCP2* (*AT5G13570*) were amplified from cDNA of wild type plants and cloned in the same vectors. For overexpression in potato, full-length cDNA sequences of SP7-like genes (including UTR regions and the protein signal peptides) were cloned into the Gateway destination vector 2xP35S-pKGWRedRoot^86^.

For yeast two hybrid assays the coding sequences of SP7-like proteins lacking the signal peptide (*RiSP7*ΔSP, *RiSP2*ΔSP, *RiSP5*ΔSP, *RiSP31*ΔSP, and GmSP1ΔSP and GrSP1ΔSP) and from *MtSR45*/*AtSR45/NbSR45*, *MtU1-70K*, *MtU2AF35b* and RiSP7 or MtSR45 truncation constructs were amplified with restriction sites and subsequently cloned into the pGBKT7 or pGADT7-REC vectors (Takara Clontech Bio Europe, Saint-Germain-en-Laye, France). The prey vector constructs containing MtSR45 and MtU2AF35b were previously reported^12,14^. For localization or co-lozalization assays in *N. benthamiana,* the coding sequences of *RiSP7*ΔSP, *RiSP2*ΔSP, *RiSP5*ΔSP, *GmSP1ΔSP*, *GrSP1ΔSP*, *MtSR45, AtSR45*, *NbSR45*, *MtU2AF35b*, *MtU1-70K*, *AtDCP2* and *RiSP7* or *MtSR45* truncation constructs were cloned into the Gateway destination vectors pCGFP-RR, pNGFP-RR, pK7WGF2 or pK7RWG2^85,87,88^. The MtSR45 mRFP co-localization construct was previously reported^14^. As co-localization control, free eGFP with stop codon was cloned into the vector pK7FWG2^87^. For *in planta* pulldown assays the coding sequences of *RiSP7*ΔSP and *RiSP7*+SP were cloned in pCGFP-RR^85^. *RiSP7*ΔSP, *RiSP2*ΔSP, *RiSP5*ΔSP, *GmSP1ΔSP*, *GrSP1ΔSP*, *MtU2AF35*, *MtU1-70K*, *RiSP7* repeat truncations as well as *MtSR45* full length or truncated cDNA versions were cloned into Gateway BiFC vectors P35S-pSPYNE-GW or P35S-pSPYCE-GW^89^ for *in planta* interaction assays.

### RNA-seq and splicing analysis

Root RNA from 6 potato plants overexpressing RiSP7 (Two biological replicates of three independent lines) and 4 empty vector control (EV) plants (Two biological replicates of 2 independent lines) was isolated following the TRIzol method. The DNaseI-treated RNA was sent to BGI (Yantian District, Shenzhen 518083, China, https://www.bgi.com/global) for sequencing on a DNBseq platform (DNA nanoballs paired-end mode based DNBSEQ Technology). An Agilent 2100 Bio analyzer (Agilent RNA 6000 Nano Kit) (Agilent Technologies Inc., United States) was used for RNA quality control. For library preparation, mRNA enrichment was realized using oligo dT beads. RNA fragmentation and first-strand cDNAs were generated using random N6-primed reverse transcription, followed by second-strand cDNA synthesis. The synthesized cDNA was subjected to end-repair, 3’-adenylation and adaptor ligation. The purified cDNA was enriched in several rounds of PCR amplifications. After sequencing, raw reads mapped to rRNA as well as low-quality reads, adaptor containing reads and reads with high content of unknown bases were removed using the BGI-internal software SOAPnuke (version 1.5.2). Clean reads were further mapped to the reference *Solanum tuberosum* v6.1^90^ genome from Phytozome using HISAT2 v2.0.4^91^ with following parameters: --phred33 --sensitive --no-discordant --no-mixed -I 1 -X 1000 --rna-strandness RF. After mapping, StringTie^92^ was used to reconstruct transcripts and novel transcripts were predicted by using Cuffcompare^93^. The coding ability of those new transcripts was assessed using CPC^94^. Novel transcripts were merged with reference transcripts using Bowtie2 v2.2.5^95^ to get a complete reference for downstream gene expression analysis. Gene expression levels for each sample were calculated with RSEM v1.2.12^96^ using default parameters and based on those the differentially expressed genes (DEGs) were detected with the DEseq2 algorithms, performed as described^97^. Genes were considered as regulated with|log2FoldChange|>1 and *pValue* < 0.05 as thresholds. For the splicing analysis, after the genome mapping, rMATS v4.0.2^98^ was used to detect genes that are differentially spliced between the groups of samples based on the relative abundances of the splicing isoforms of a gene across samples with following parameters: -t paired --nthread 8 --tstat 4. It calculated the *pValue* and false discovery rate (FDR) for the difference in the isoform ratio of genes in the EV samples versus the SP7-overexpressing samples. Genes with FDR ≤ 0.05 were defined as differential alternative splicing (DAS) genes (Supplementary Table 4).

### Gene expression analyses

Relative expression values for the specific mRNA isoforms alternatively spliced detected in Supplementary Table 4 were verified by qRT-PCR. To that end cDNA synthesis and quantitative real time PCR (qPCR) were carried out as described^86^. The PCR protocol was selected as follows: 5 min 95 °C, 15 s 95 °C, 15 s Tm °C, 30 s 72 °C (40 cycles) on a CFX Connect Real-Time PCR Detection System (Bio-Rad Laboratories GmbH, Munich, Germany). The annealing temperature Tm was adjusted for each primer combination. For normalization of plant and fungal transcript levels the actin gene of potato *StActin* (Soltu.DM.05G024990), was used. Expression data are given as relative expression values calculated using the 2^-ΔCt^ method. Transcript levels of genes were determined in 3 independent lines ectopically expressing *RiSP7*, 1 line expressing *RiSP5* and 2 control lines expressing an empty vector (EV) in at least three biological replicates with two technical replicates per reaction. All primers are listed in Supplementary Table 5.

### Yeast two hybrid interaction assays

pGBKT7 bait vectors were transformed in *S. cerevisiae* AH109 (Takara Clontech Bio Europe, Saint-Germain-en-Laye, France) using the LiAc/SS carrier DNA/PEG Method^99^. Yeast-two-hybrid library screens were carried out according to Matchmaker® Gold Yeast Two-Hybrid System User Manual (Takara Clontech Bio Europe, PT4084-1) using a cDNA library of *M. truncatula* roots colonised with *R. irregularis* in *S. cerevisiae* strain Y187^12^ or a normalized cDNA library of *A. thaliana* (Takara Clontech Bio Europe, Cat.# 630487). For direct interaction studies co-transformation of bait and pray was employed. Dilution series were spotted on yeast synthetic drop-out medium without leucine and tryptophan and on synthetic drop-out medium without leucine, tryptophan, histidine and adenine under sterile conditions and incubated for 4-5 days at 30°C. Positive interactions are indicated by yeast colony growth on media lacking leucine, tryptophan, histidine and adenine.

### Transient protein expression in *N. benthamiana* leaves

Transient expression of proteins in *N. benthamiana* leaves was performed as described^89^ using *Agrobacterium tumefaciens* strain GV3101 to infiltrate leaves of 4-5 weeks old plants. Final optical density (OD_600_) of *A. tumefaciens* was adjusted to 0.5 with AS medium supplemented with 2% sucrose (localization and co-localization studies) or to 1.0 (GFP pulldown assay and BiFC). To prevent silencing, an *A. tumefaciens* strain K5A018 carrying the p19 silencing suppressor of tomato bushy stunt virus^100^ was co-infiltrated in a 1:1 ratio for single transformation (localization and pulldown assay) or a 1:1:1 ratio for co-transformation (co-localization or BiFC). Leaf discs were excised 2-4 dpi and fluorescence was analyzed in epidermal cells using confocal microscopy.

### Pulldown assay from *N. benthamiana* leaves and peptide identification

*N. benthamiana* leaves were transiently transformed as described above with *A. tumefaciens* (strain GV3101) carrying the vectors pCGFP-RR::SP7ΔSP, pCGFP-RR::SP7+SP and pCGFP-RR::eGFP^101^. Total protein extraction from *N. benthamiana* leaves and pulldown assay were carried out according to^102^ with the following modifications: For immunoprecipitation of GFP-tagged proteins 50 µl of GFP-Trap magnetic beads (Chromotek, Planegg-Martinsried, Germany) were washed 3 times with extraction buffer and incubated for 2 h at 4 °C with 500 µl of a total protein extract containing a protease inhibitor cocktail (1x cOmplete^™^, Mini, EDTA-free, Sigma-Aldrich, 04693159001). Proteins were eluted from magnetic beads by boiling at 95°C for 10 min in 50 µl 2x Laemmli sample buffer. For protein identification 25 µl of the eluted protein fraction were loaded on a SDS gel. Protein samples were sliced out as a compact band and were analyzed by mass spectrometry (Toplab GmbH, Martinsried, Germany). Protein samples were reduced with DTT and alkylated with IAA prior to a tryptic digestion (MS grade, Serva) over-night at 37°C. For nano-ESILC-MS/MS analysis an aliquot of the peptide solution was used. HPLC separation was done using an EASY-nLC1000 (Thermo Scientific) system with the following columns and chromatographic settings: The peptides were applied to a C18 column (Acclaim® PepMap 100 pre-column, C18, 3 μm, 2 cm x 75 μm Nanoviper, Thermo Scientific) and subsequently separated using an analytical column (EASY-Spray column, 50 cm x 75 μm ID, PepMap C18 2 μm particles, 100 Å pore size, Thermo Scientific) by applying a linear gradient (A: 0.1% formic acid in water, B: 0.1% formic acid in 84% ACN) at a flow rate of 200 nl/min. The gradient used was: 1-30% B in 80 minutes, 30-60% B in 20 minutes, 100% B 10 minutes. Mass-spectrometric analysis was done on a LTQ Orbitrap XL mass-spectrometer (Thermo Scientific), which was coupled to the HPLC-system. The mass-spectrometer was operated in the so-called “data-dependent” mode where after each global scan, the five most intense peptide signals are chosen automatically for MS/MS-analysis. The detected peptide masses were compared with all sequences from *Nicotiana benthamiana* draft genome sequence v1.0.1 at the Solanaceae Genomics Network database (https://solgenomics.net/). The identification threshold was set to p <0.05. Carbamidomethylation of cysteines was set as fixed modification and oxidized methionine as variable. Only peptides with an Ion score >30 were considered as significant. Proteins identified in the free eGFP samples were subtracted as background. Gene descriptions for identified *Arabidopsis*orthologs were retrieved from TAIR (https://www.arabidopsis.org/tools/bulk/genes/index.jsp).

### Confocal microscopy and image processing

Confocal microscopy images were taken using a Leica TCS SP5 (DM5000) and the LASAF v2.6 software. A HCX PL APO CS 20.0× (NA 0.70) DRY UV (Leica, Wetzlar, Germany) was used at 21°C. In localization and co-localization studies the fluorescence proteins eGFP (488 nm) and mRFP (561 nm) were excited with an argon and a DPSS 561 laser, respectively. Emission of eGFP was detected from 493 – 530/550 nm and mRFP from 566 – 670/726 nm. To better discriminate between chloroplast autofluorescence and nuclear bodies in the DsRed and mRFP channels, emissions in far red (675-765 nm) were additionally captured in a separate channel and pseudo coloured magenta in merged pictures when necessary. For BiFC analyses YFP emission was detected from 525 - 590 nm after excitation at 514 nm. Pictures were processed using Fiji, https://imagej.net/software/fiji/downloads^103^.

### Data availability

The sequences of the SP7-like effectors are available in GenBank with the following accessions: MF521604 (*RiSP7*), MF521605 (*RiSP2*), MF521606 (*RiSP5*), MF521607 (*RiSP31*), MF521608 (*GmarSP1*), MF521609 (*GrosSP1*). *M. truncatula* gene names are from the genome release (Mt4.0v2, http://www.medicagogenome.org/about/project-overview).

### Statistical analysis

Gene expression data are shown as boxplots in which the three horizontal lines represent the 3 quartiles, the “x” represents the average and the whiskers reach out to the minimum and maximum values, except for values deemed as outliers, which are shown as black dots. The Shapiro-Wilk test was carried out to check the normal distribution of the data and the Levene’s test was used to check the homoscedasticity between each two groups compared. When two groups compared were normally distributed, a T-test was carried out in Microsoft Excel applying the corresponding correction depending on whether the homoscedasticity could be assumed or not. When one or both groups were not normally distributed, they were compared with a Mann-Whitney U test. Significance levels are shown as follows: “ns” (non-significant, p > 0.05), “*” (significant p < 0.05), “**” (significant p < 0.01) or “***” (significant p < 0.001). Except for the T-tests, all statistical tests were carried out on the website Statistics Kingdom (www.statskingdom.com) and the box plots were generated with the RStudio software^104^ using the package ggplot2.

## Data availability

Source data are provided with this paper.

## Acknowledgements

We thank Dr. J. Palenzuela (EEZ, CSIC, Spain), Dr. C. Roux (CNRS, Toulouse, France) and Dr. S. Roy (Agronutrition, France) for the generous gift of inocula from *G. margarita* and *G. rosea*. We thank the funding support of the Future Fields KIT Project “Plants fit for Future” to N.R.

## Author contributions

R.B., S.H., D.F-G., and N.R. designed the experiments. R.B., S.H., D.F-G. performed the assays. R.B., S.H., D.F-G., T.L. and N.R. interpreted the data. R.B. and N.R. wrote the manuscript with inputs from all the authors.

## Additional Information

**Suppplementary Information.** The online version contains supplementary material available at **XXX**.

**Correspondence** and requests for materials should be addressed to Natalia Requena

## Supplementary Figures

**Supplementary Figure 1.**
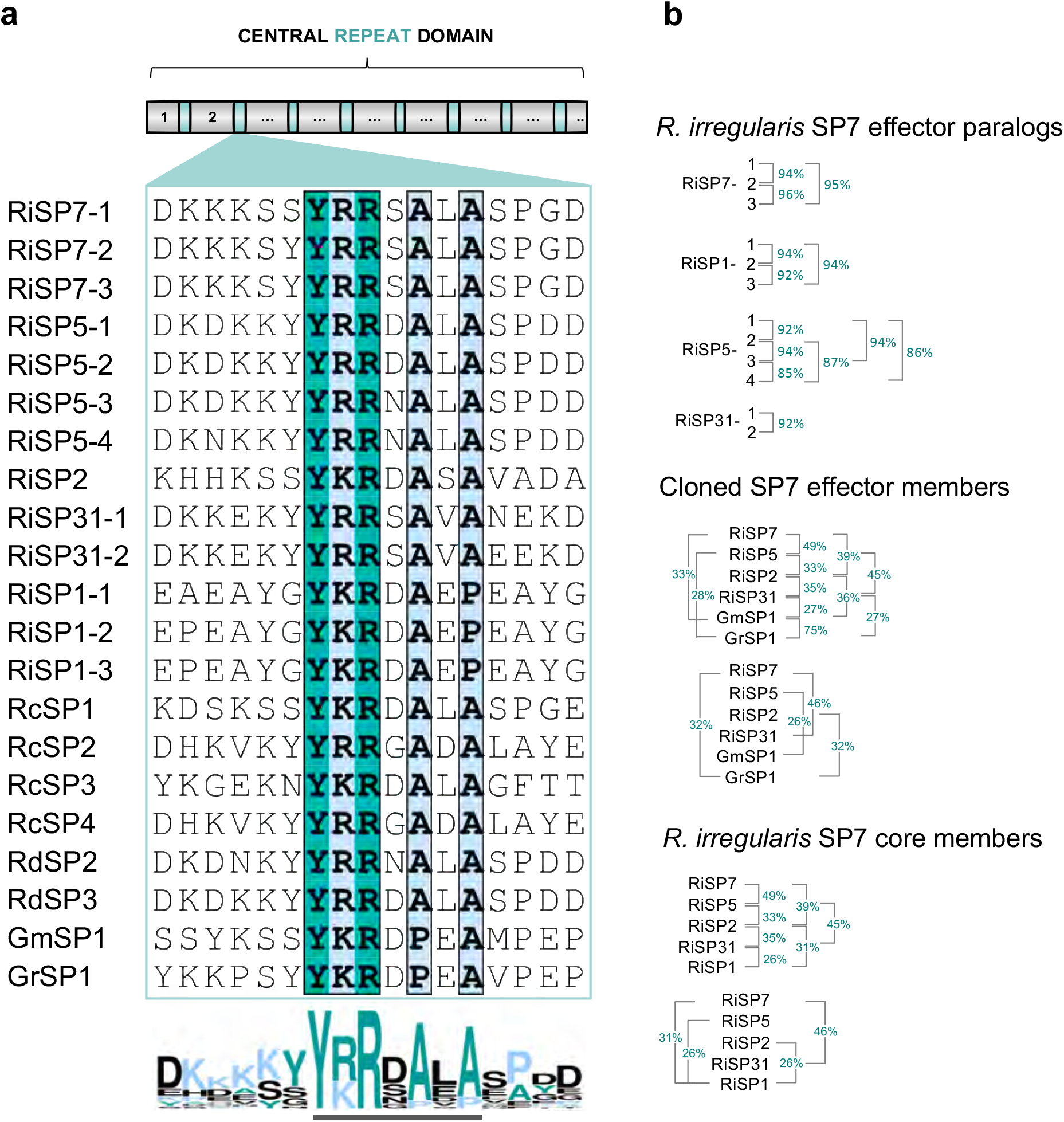
Identification of SP7 family characteristic conserved islands and effector motif. **a** All identified full length SP7-like effectors (Supplementary Table 2) contain family conserved amino acid stretches (islands) within each repeat subunits. Shown is an alignment of the conserved islands from the 2^nd^ repeat of each effector member. Ri (*Rhizophagus irregularis*), Rc (*Rhizophagus clarus*), Rd (*Rhizophagus diaphanus*), Gm (*Gigaspora margarita*), Gr (*Gigaspora rosea*). Bottom Logo represents consensus sequence (Weblogo) of the conserved islands, revealing the newly identified SP7 effector family motif Y-[KR]-R-X-[AP]-X-[AP] (underlined). Aligned sequences were visualized using ESpript3.0 (Blosom62 score). Identical amino acids are highlighted with bold letters, green background and conserved amino acids with bold letters, blue background. **b** Comparison of percentage protein identities between *R. irregularis* SP7-like paralogs, cloned and functionally analyzed SP7-like members and SP7-like core effectors present in *R. irregularis* after multiple ClustalO alignments. For protein identities between all identified members see Supplementary Table 2.

**Supplementary Figure 2.**
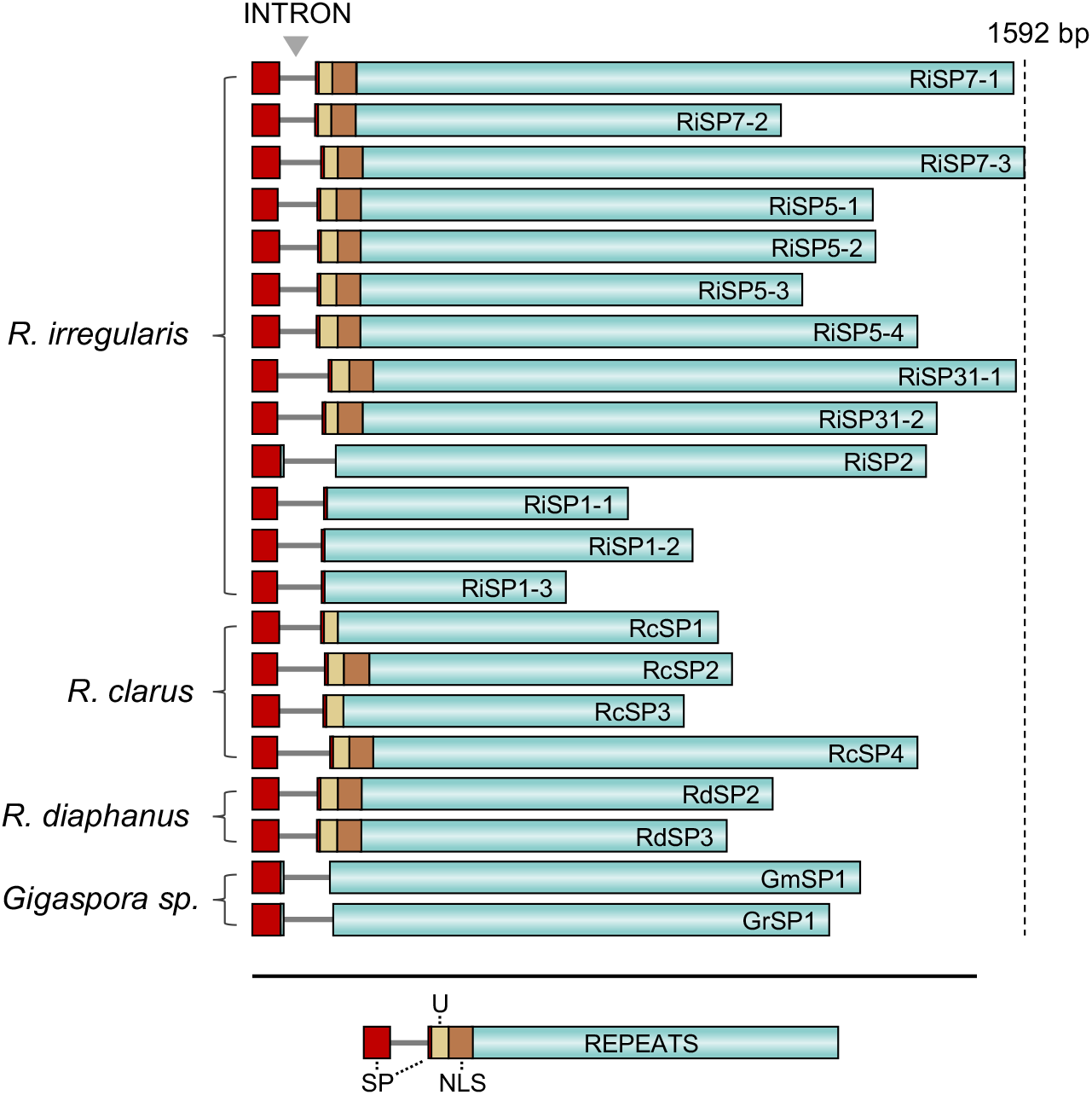
Analysis of the SP7-like effector family gene organization reveals position conserved introns. Illustrated are the coding sequences of all identified full lenght SP7 family effectors (Supplementary Table 2) from four different Glomeromycotina fungal species. All family members contain a single intron at the N-terminal region prior to the central repeat regions. For most effectors, the intron disrupts the SP coding sequences. Signal peptide (SP) highlighted in red, conserved domain with unknown function (U) in yellow, NLS in brown and repeat regions in green. Gene illustrations are drawn to scale.

**Supplementary Figure 3.**
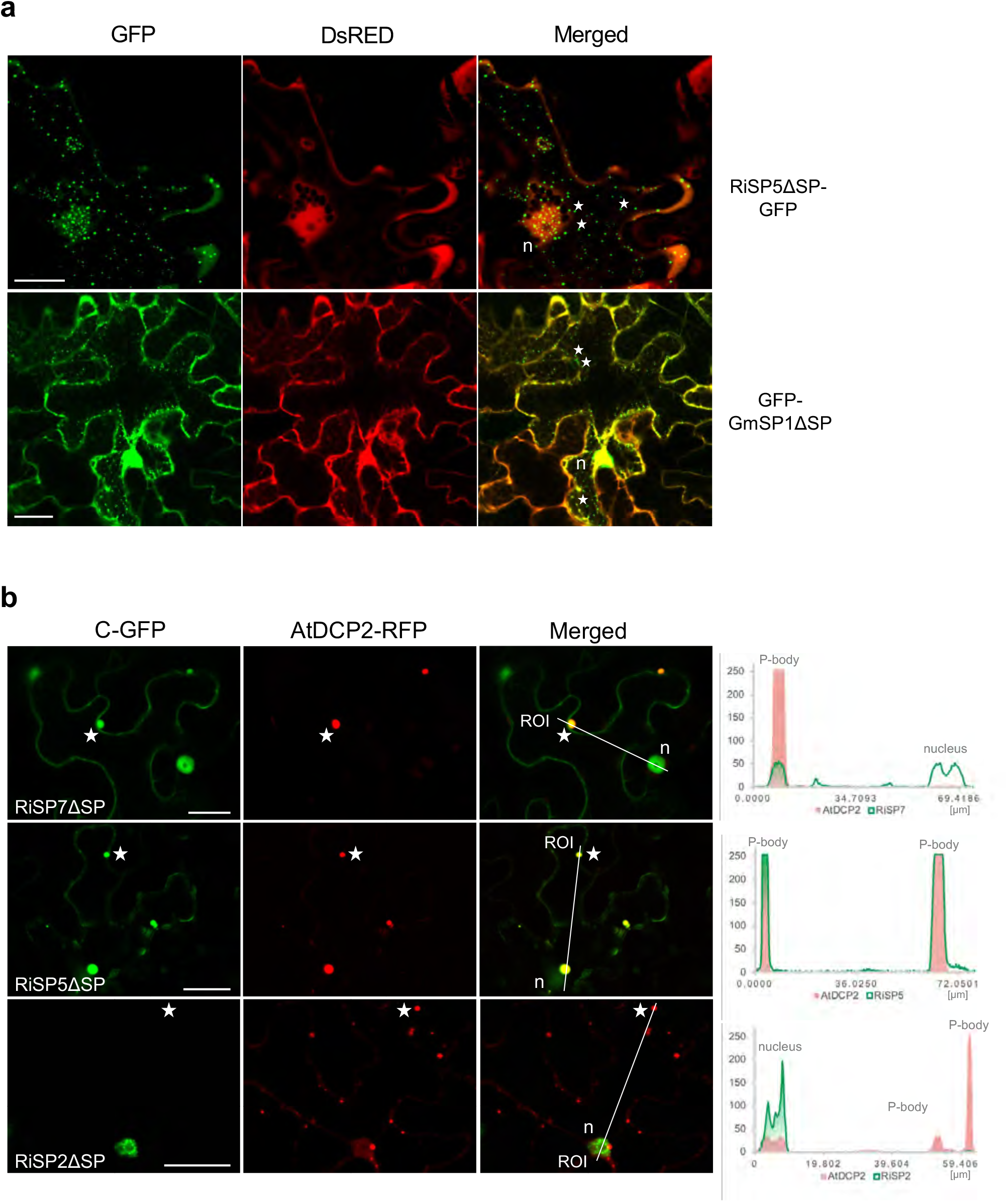
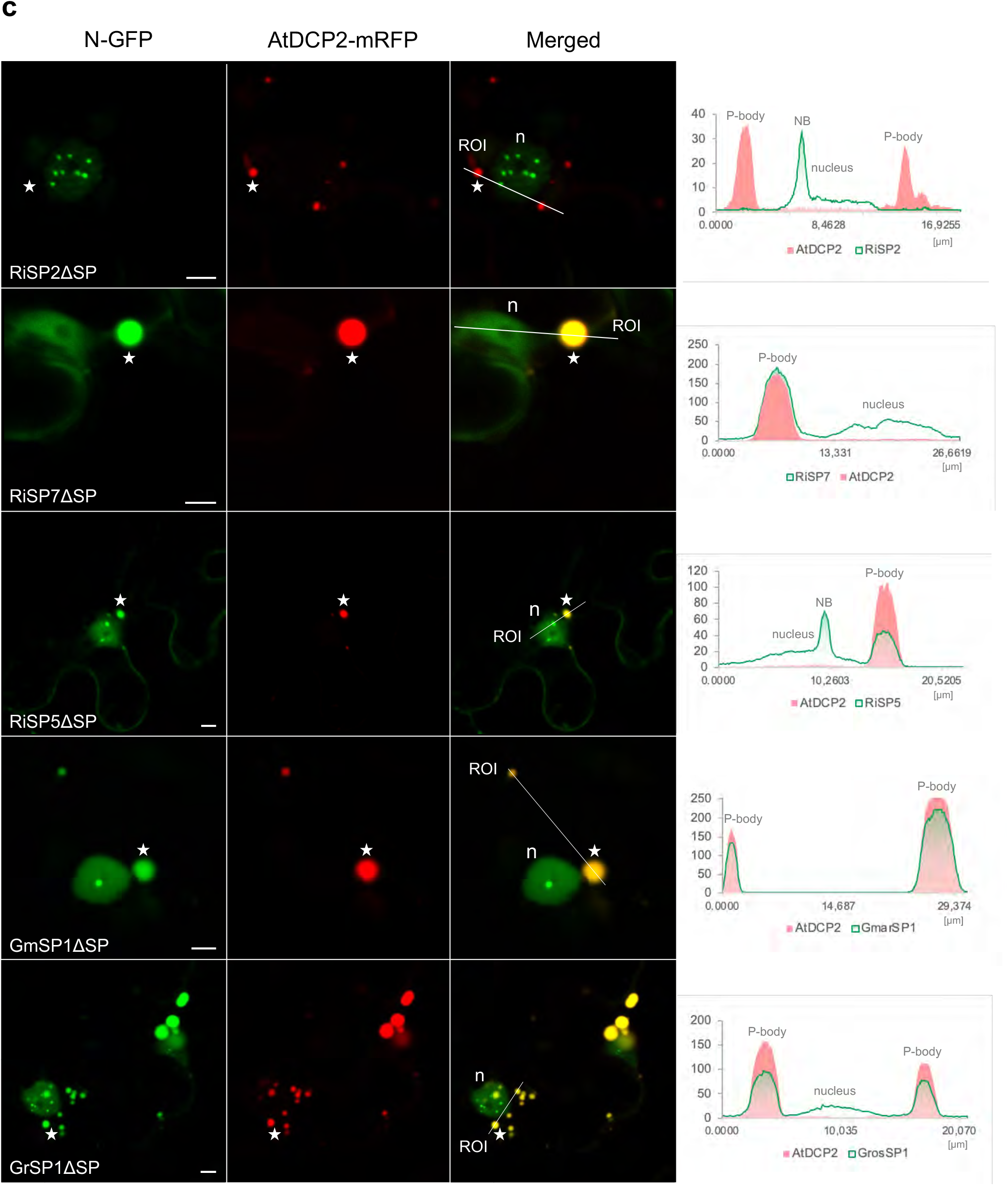
Co-localization study of SP7 family members with AtDCP2 P-body marker in *N. benthamiana*. **a** Example pictures of cytoplasmic condensates occasionally found in a few cells for some SP7-like effectors. Shown are RiSP5ΔSP with C-terminal GFP fusion and GmSP1ΔSP with N-terminal GFP fusion expressed together with free DsRed as control to visualize nucleus and cytoplasm in *N. benthamiana* epidermal cells. Cytoplasmic condensates were absent from the DsRed channel. Exemplary cytoplasmic condensates for RiSP5ΔSP and GmSP1ΔSP are marked with stars. Scale bars represent 25 µm; n = nucleus. **b-c** Co-localization of SP7-like effectors without SP and fused to either C-terminal GFP (b, C-GFP) or N-terminal GFP (c, N-GFP) together with P-body marker AtDCP2 C-terminally fused to mRFP in *N. benthamiana* epidermal cells. With exception of nuclear localized RiSP2ΔSP, all tested effectors localized to the same cytoplasmic P-bodies marked by AtDCP2. Exemplary P-bodies are marked with stars. Fluorescence intensity blots generated from individual GFP and mRFP channels along transection lines are shown next to each panel. At positions of mRFP marked P-bodies, increase of GFP fluorescence can be observed for all SP7-like proteins but RiSP2. Scale bars represent 25 µm (b) and 5 µm (c). ROI = start of intensity measurement. NB = nuclear body. n = nucleus.

**Supplementary Figure 4.**
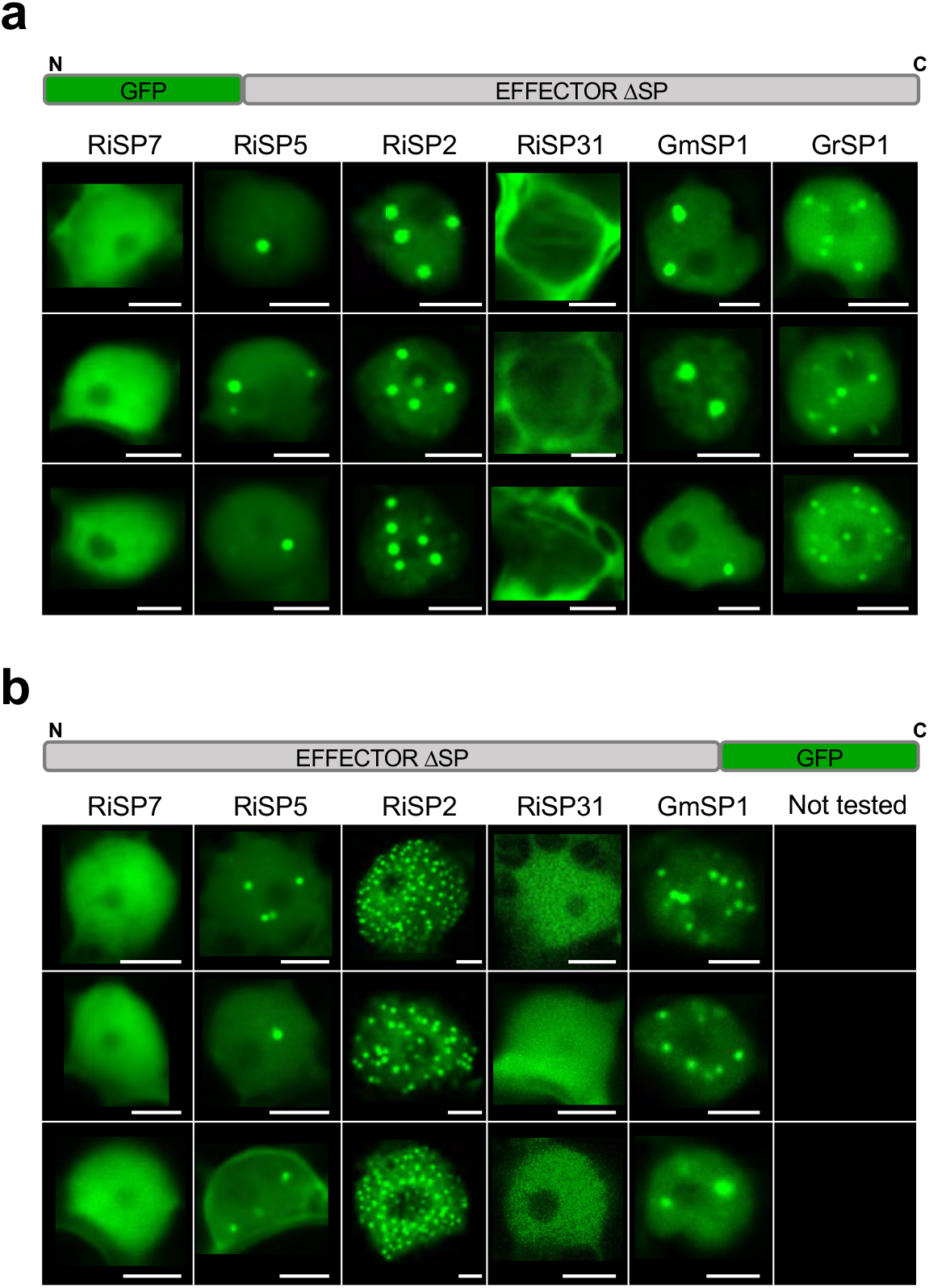
Nuclear localization patterns of SP7-like effectors fused N- or C-terminally to eGFP in *Nicotiana benthamiana*. **a-b** Shown are always three representative plant nuclei with the localization pattern of SP7-like effector proteins lacking their signal peptide (ΔSP) fused either N- (a) or C-terminally (b) to eGFP after transient expression in *N. benthamiana* epidermal cells. RiSP2, RiSP5, GmSP1 and GrSP1 localize to the plant nucleus and accumulate in nuclear bodies. Number and sizes of nuclear bodies differ, ranging from one to many bodies. RiSP7 N- and C-terminal GFP constructs lack the formation of nuclear bodies and show evenly distributed fluorescent signals throughout the nucleoplasm instead. GFP-RiSP31 is strongly located in the nuclear surroundings often with visible transversally stripes spanning the nucleus, while RiSP31-GFP can only be weakly detected in diffuse signals within nuclei as indicated by its grain-like signals. Scale bars represent 5 µm.

**Supplementary Figure 5.**
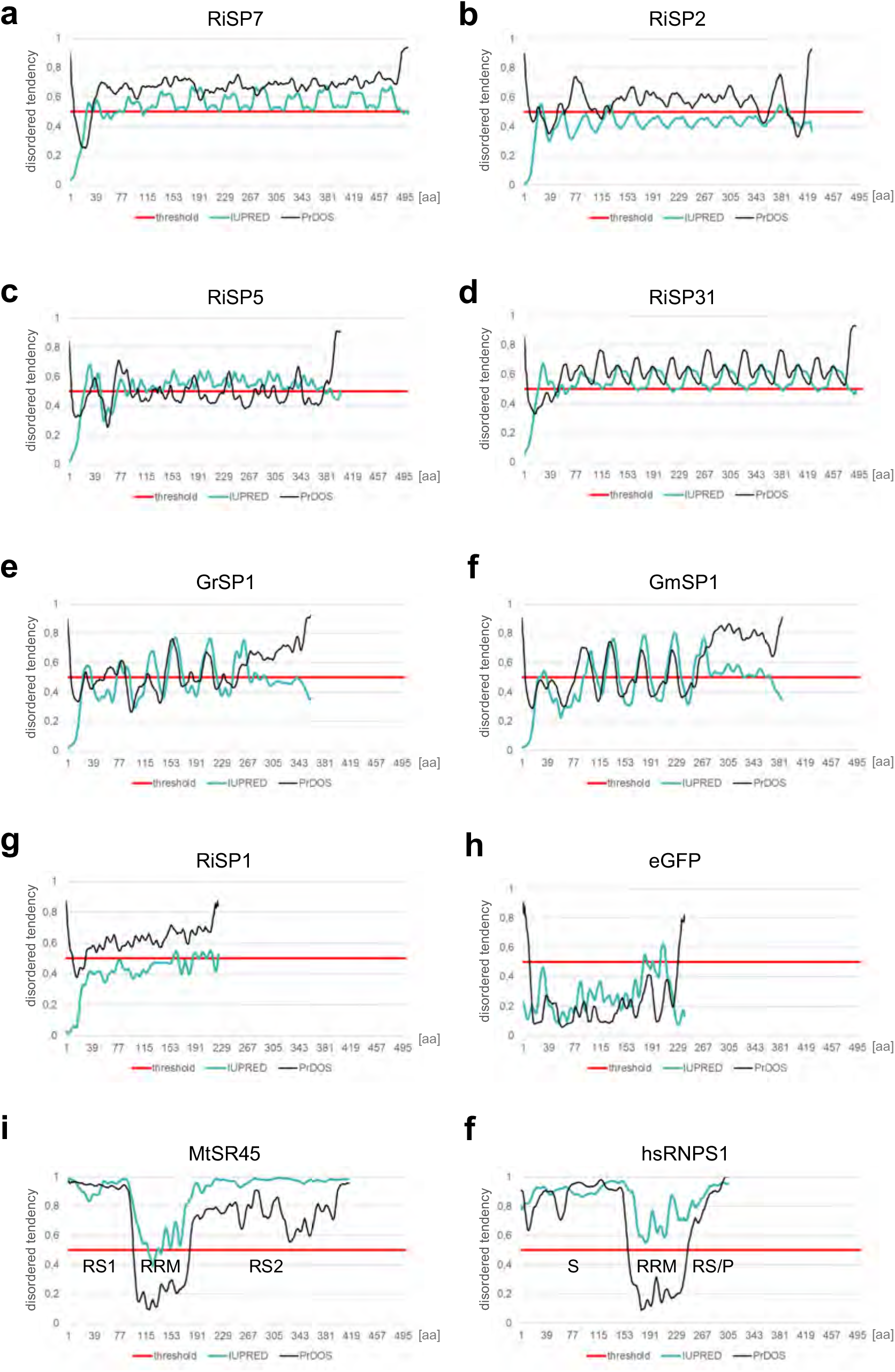
Disorder probabilities of SP7-like effectors analyzed by PrDOS and IUPred prediction softwares. **a-g** All SP7-like proteins showed an overall disordered tendency. **h** eGFP is, in contrast, structurally ordered. **i-f** MtSR45 and hsRNPS1 also show a disordered tendency in agreement with the predicted disordered RS domains of SR proteins (Haynes and Iakoucheva, 2006), while the RRM domains are less disordered. Threshold for disorder probability was 0.5 marked with a red line. False positive rate (PrDOS) 5%.

---

**Supplementary Figure 6.**
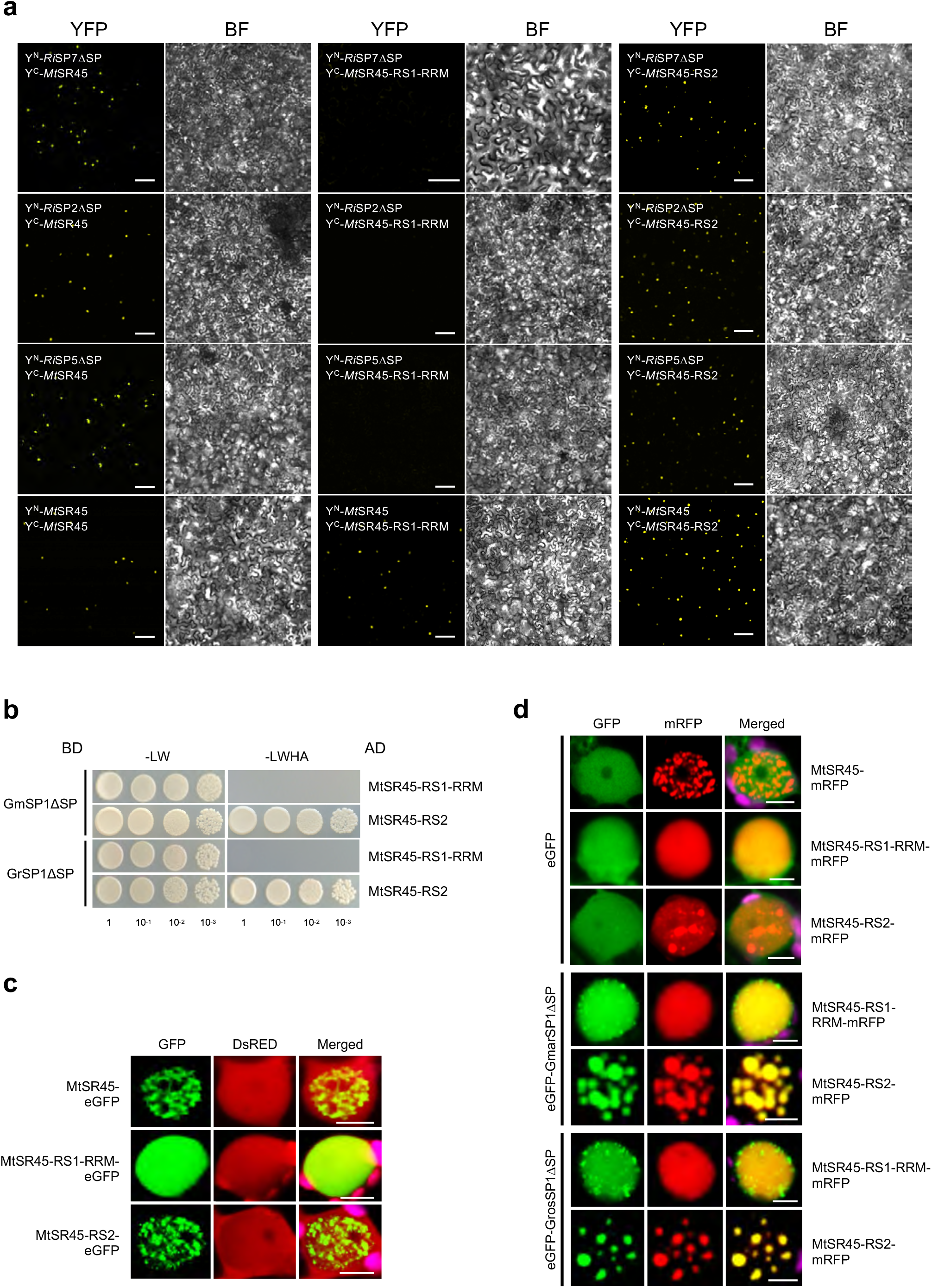
Additional interaction and localization assays of MtSR45 truncations and SP7-like effectors. **a** Overview pictures for BiFC assays between RiSP7ΔSP, RiSP2ΔSP or RiSP5ΔSP (fused to N-terminal half of YFP, YFP^N^). and MtSR45 or the truncation domains RS1-RRM (MtSR45-RS1-RRM) and RS2 (MtSR45-RS2) fused to C-terminal half of YFP, YFP^C^. Shown are pictures of infiltrated *N. benthamiana* whole cell tissue areas. Reconstituted YFP signals (left panels) in nuclei of several cells demonstrated interactions between SP7-like effectors and full length MtSR45 as well as the RS2 domain alone. No YFP signals were detected for RS1-RRM truncations. In addition, interaction of MtSR45 with itself as well as with all MtSR45 truncations was observed. Right panels: Brightfield pictures (BF) of respective cell areas seen in YFP channels. Scale bars represent 100 µm. **b** The RS2 domain of MtSR45 is sufficient and required for interaction with *Gigaspora* SP7-like effectors. Shown are Y2H interaction spotting assays (using yeast dilution series 1-10^-3^) between GmSP1ΔSP or GrSP1ΔSP as bait (BD) and the RS1-RRM or RS2 truncations as prey (AD). Positive interactions are indicated by colony growth on media lacking leucine, tryptophan, histidine, adenine (-LWHA). EV = Empty vector. **c-d** single plant nuclei are shown after transient expression of fluorescent tagged proteins in *N. benthamiana* epidermal cells. Chloroplast autofluorescences are pseudo-coloured in magenta (merged pictures). Scale bars represent 5 µm. **c** Subcellular localization of MtSR45 and truncations (with C-terminal eGFP tag). Only full length MtSR45 and the RS2 domain localized to nuclear speckles. As control, free DsRed was co-expressed. **d** Co-localization assay of GmSP1ΔSP and GrSP1ΔSP (N-terminal eGFP tag) with MtSR45, RS1-RRM or RS2 truncation domains (C-terminal mRFP tag). Both *Gigaspora* effectors only co-localized with full length MtSR45 and the RS2 domain in nuclear bodies. All experiments were performed with SP7-like effectors lacking their signal peptides (ΔSP).

**Supplementary Figure 7.**
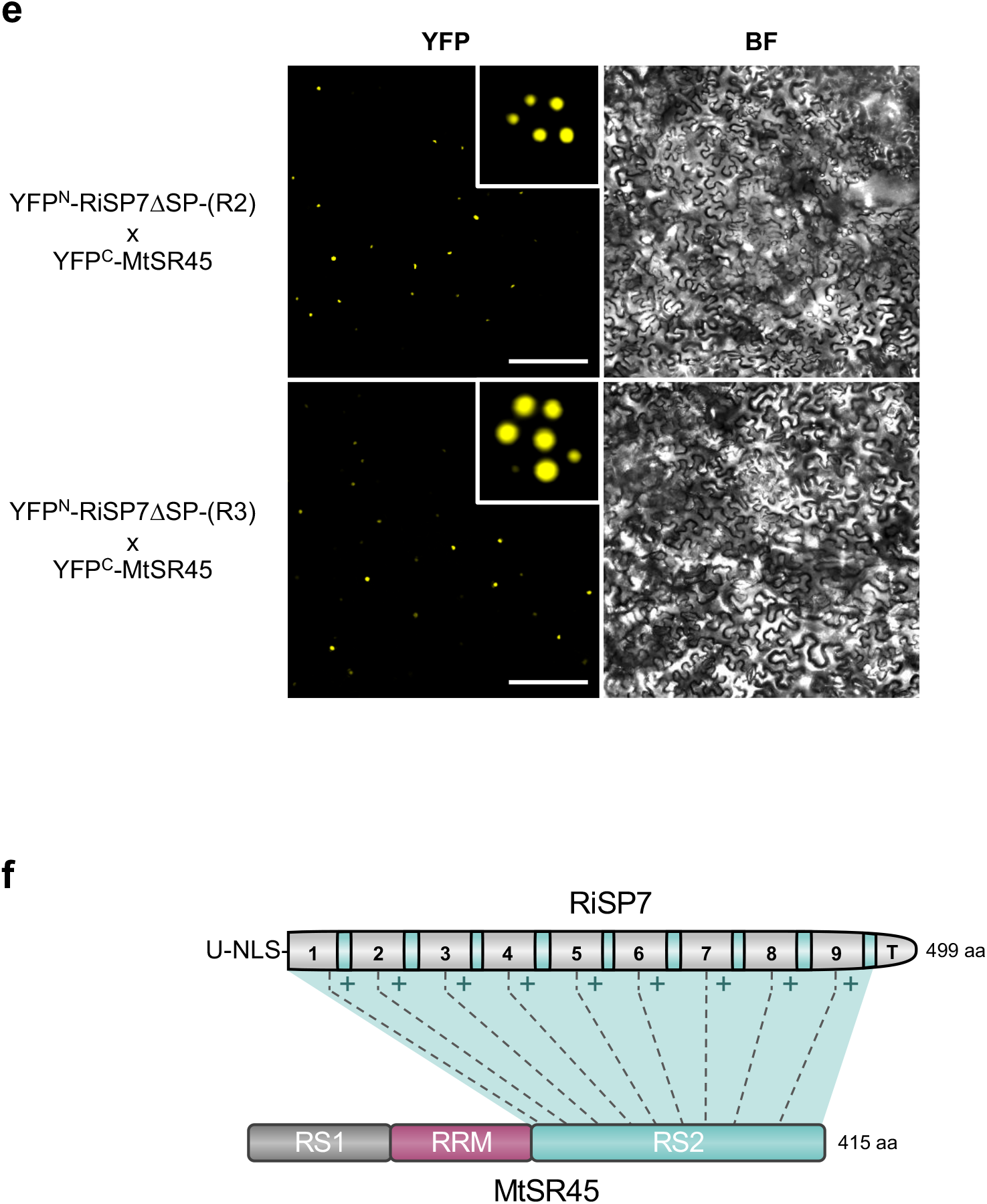
Number of RiSP7 repeats determines its interaction strength with MtSR45. **a** Overview pictures of BiFC assays using RiSP7ΔSP truncations (without SP, fused to N-terminal half of YFP, YFP^N^) together with MtSR45 (fused to C-terminal half of YFP, YFP^C^) after transient expression in *N. benthamiana* epidermal cells. Shown are pictures of infiltrated *N. benthamiana* whole cell tissue areas. Reconstituted YFP signals in nuclear condensates could be observed for truncations with two and three repeat units in several cell nuclei (inlet pictures show exemplary magnified single nuclei with nuclear bodies). BF = Brightfield pictures of respective cell areas seen in YFP channels. Scale bars represent 200 µm. **b** Model of proposed mode of interaction between RiSP7 and the RS2 domain of SR45. Each single repeat unit enhances the affinity and interaction strength (+) with the RS2 domain.

**Supplementary Figure 8.**
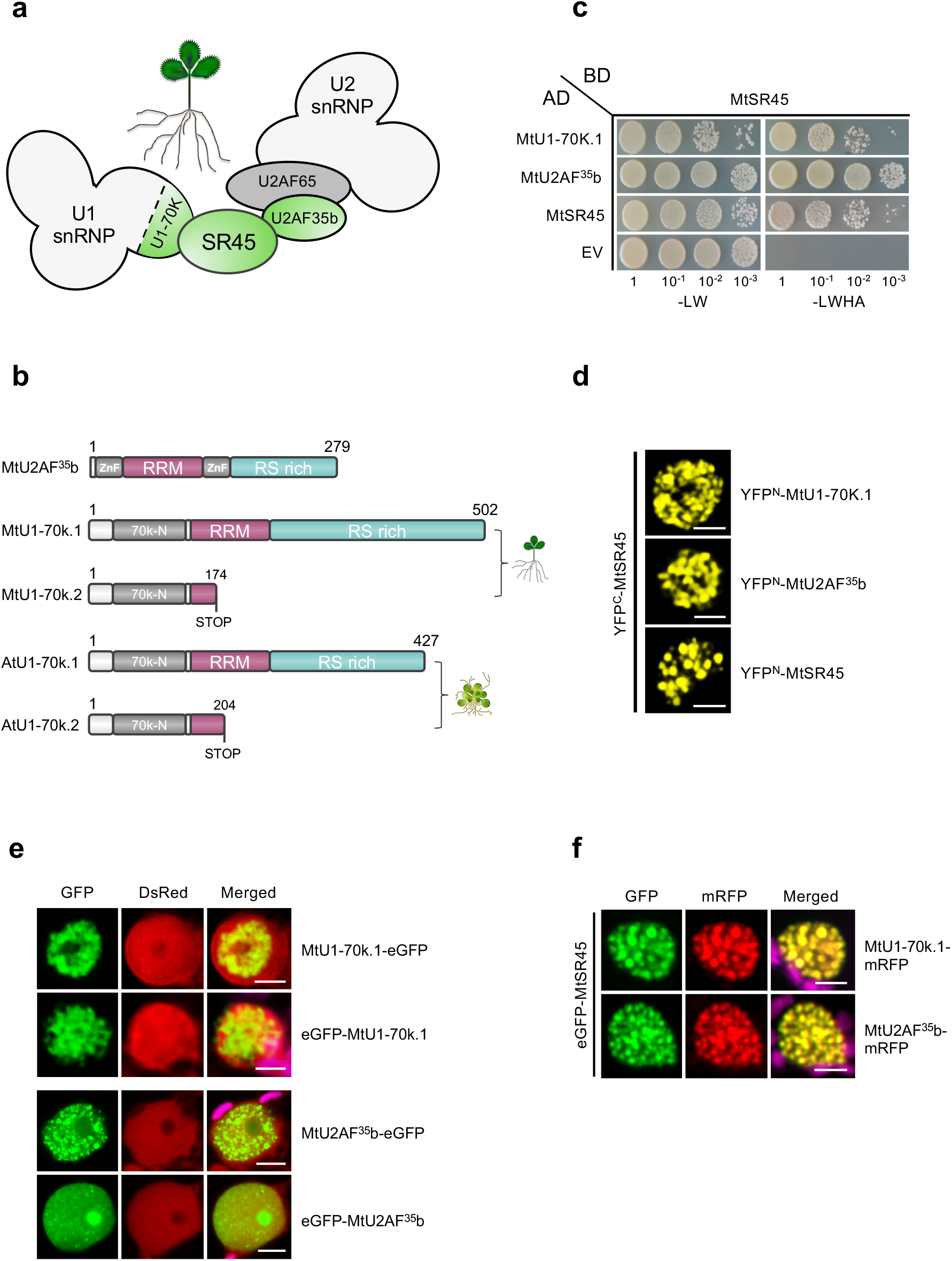
SR45 interactions with spliceosomal components U2AF^35^b and U1-70k are conserved in *Medicago*. **a** Model for possible interaction network of *Medicago* SR45 with MtU2AF^35^b and MtU1-70k based on published interactions of *Arabidopsis* SR45 (see Fig. 6). **b** Protein structures of MtU2AF^35^b and MtU1-70k. MtU2AF^35^b comprises a RRM motif flanked by two CCCH-type zinc fingers (ZnFs) and a RS-rich domain similar to its *Arabidopsis* orthologue (Wang & Brendel, 2006). MtU1-70K exists as two isoforms (MtU1-70k.1 and MtU1-70k.2) also described for the AtU1-70k ortholog (Golovkin & Reddy, 1996) with a conserved N-terminal domain (70K-N). Isoform MtU1-70k.2 lacks RRM and RS-rich domains. **c** Y2H assays revealed the interaction between MtSR45 (Bait, BD) with *Medicago* U2AF^35^b and U1-70k.1 (Preys, AD). MtSR45 was also tested for homo-dimerization as previously reported for plant SR proteins (Golovkin & Reddy, 1999; Tanabe *et al*., 2009). Positive interactions are indicated by yeast colony growth on media lacking leucine, tryptophan, histidine, adenine (-LWHA) using dilution series (1-10^-3^). EV = Empty vector. **d-f** Shown are localization patterns in single plant nuclei after transient expression of fluorescent tagged proteins in *N. benthamiana* epidermal cells. Scale bars represent 5 µm. **e-f** Chloroplast autofluorescences are pseudo-coloured in magenta (merged pictures). **d** Confirmation of interactions seen in (c) using BiFC. Interaction was detected in nuclear speckles for MtSR45 (fused to C-terminal half of YFP, YFP^C^) co-expressed with either MtU2AF^35^b, MtU1-70k.1 or with itself (fused to N-terminal half of YFP, YFP^N^). **e** Subcelluar localization of MtU1-70k.1 or MtU2AF^35^b (N- or C-terminal eGFP fusions) showed localizations in differently shaped nuclear speckles. Additionally, eGFP-MtU2AF^35^b localized in the nucleolus. As control, free DsRed was co-expressed. **f** Co-localization of eGFP-MtSR45 with MtU1-70k.1 or MtU2AF^35^b fused C-terminally to mRFP. Both proteins co-localized with MtSR45 in nuclear speckles.

**Supplementary Figure 9.**
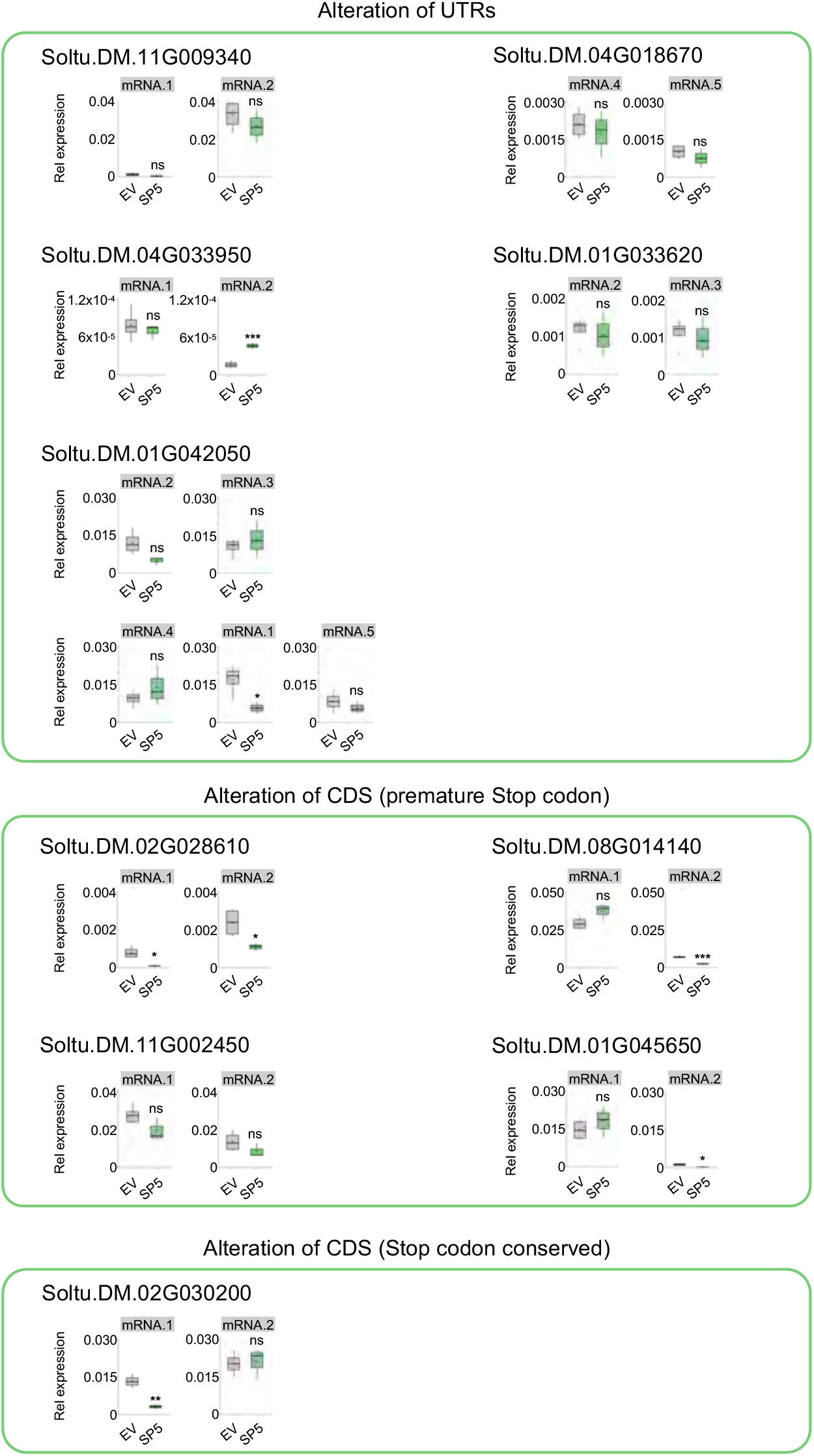
qRT-PCR analysis of the splicing in response to ectopic expression of RiSP5 in potato. qRT-PCR quantification of the relative expression of each splicing isoform of the selected DAS potato genes in roots of control plants (EV) and plants ectopically expression RiSP5. Statistical significance was assessed by performing a Student T test or a Mann Whitney U test after checking the normal distribution of the data and homocedasticity as explained in Materials and Methods. Significance levels: n.s. (non-significant) p>0.05, * p<0.05, ** p<0.01, *** p<0.001.

**Supplementary Figure 10.**
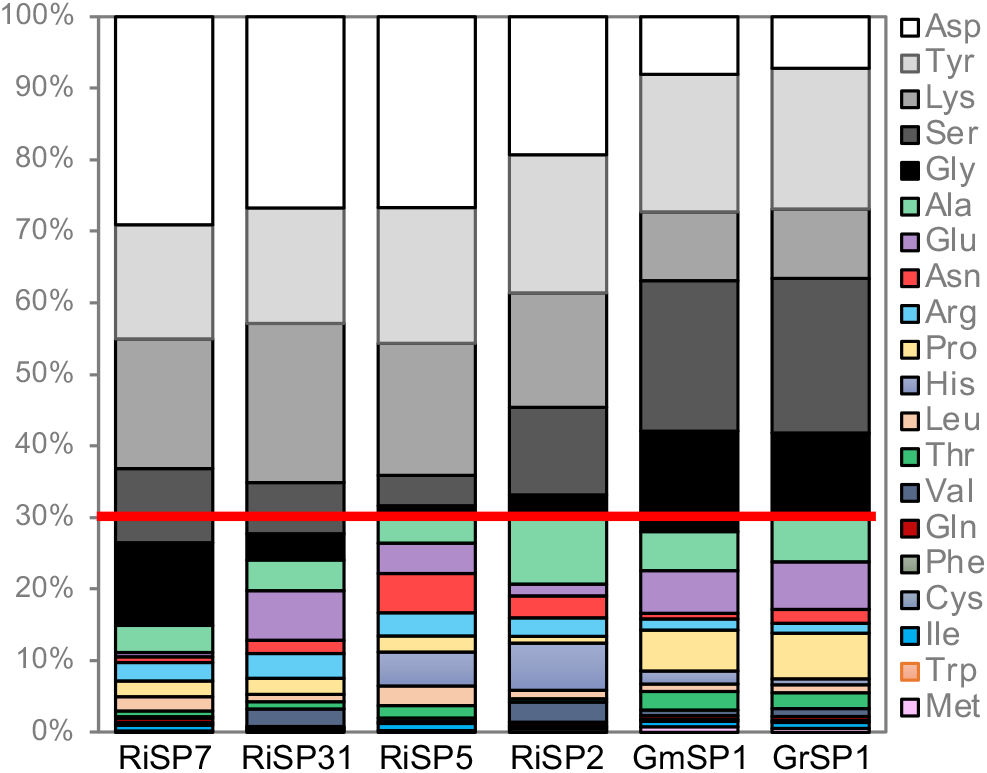
Relative aa composition for each SP7 family member. The majority of each effector (between 70-85 %) of each effector is composed of only 5 different aa (Asp, Tyr, Lys, Ser; white-black colors). Red line indicates 70% proportion.

## References

1 Chisholm, S. T., Coaker, G., Day, B. & Staskawicz, B. J. Host-microbe interactions: shaping the evolution of the plant immune response. Cell 124, 803–814 (2006). 10.1016/j.cell.2006.02.008

2 Weiberg, A., Wang, M., Bellinger, M. & Jin, H. Small RNAs: a new paradigm in plant-microbe interactions. Annu Rev Phytopathol 52, 495–516 (2014). 10.1146/annurev-phyto-102313-045933

3 Maroti, G., Downie, J. A. & Kondorosi, E. Plant cysteine-rich peptides that inhibit pathogen growth and control rhizobial differentiation in legume nodules. Curr Opin Plant Biol 26, 57–63 (2015). 10.1016/j.pbi.2015.05.031

4 Yu, H. et al. Plant lysin motif extracellular proteins are required for arbuscular mycorrhizal symbiosis. Proc Natl Acad Sci U S A 120, e2301884120 (2023). 10.1073/pnas.2301884120

5 Ratu, S. T. N., Amelia, L. & Okazaki, S. Type III effector provides a novel symbiotic pathway in legume-rhizobia symbiosis. Biosci Biotechnol Biochem 87, 28–37 (2022). 10.1093/bbb/zbac178

6 Khan, A. et al. Effector-triggered inhibition of nodulation: A rhizobial effector protease targets soybean kinase GmPBS1-1. Plant Physiol 189, 2382–2395 (2022). 10.1093/plphys/kiac205

7 Tisserant, E. et al. Genome of an arbuscular mycorrhizal fungus provides insight into the oldest plant symbiosis. Proc Natl Acad Sci U S A 110, 20117–20122 (2013). 10.1073/pnas.1313452110

8 Lin, K. et al. Single nucleus genome sequencing reveals high similarity among nuclei of an endomycorrhizal fungus. PLoS Genet 10, e1004078 (2014). 10.1371/journal.pgen.1004078

9 Yildirir, G. et al. Long reads and Hi-C sequencing illuminate the two-compartment genome of the model arbuscular mycorrhizal symbiont Rhizophagus irregularis. New Phytol 233, 1097–1107 (2022). 10.1111/nph.17842

10 Manley, B. F. et al. A highly contiguous genome assembly reveals sources of genomic novelty in the symbiotic fungus Rhizophagus irregularis. G3 (Bethesda) 13 (2023). 10.1093/g3journal/jkad077

11 Teulet, A. et al. A pathogen effector FOLD diversified in symbiotic fungi. New Phytol 239, 1127–1139 (2023). 10.1111/nph.18996

12 Kloppholz, S., Kuhn, H. & Requena, N. A secreted fungal effector of Glomus intraradices promotes symbiotic biotrophy. Curr Biol 21, 1204–1209 (2011). 10.1016/j.cub.2011.06.044

13 Tsuzuki, S., Handa, Y., Takeda, N. & Kawaguchi, M. Strigolactone-Induced Putative Secreted Protein 1 Is Required for the Establishment of Symbiosis by the Arbuscular Mycorrhizal Fungus Rhizophagus irregularis. Mol Plant Microbe Interact 29, 277–286 (2016). 10.1094/MPMI-10-15-0234-R

14 Voß, S., Betz, R., Heidt, S., Corradi, N. & Requena, N. RiCRN1, a Crinkler Effector From the Arbuscular Mycorrhizal Fungus Rhizophagus irregularis, Functions in Arbuscule Development. Frontiers in Microbiology 9 (2018). 10.3389/fmicb.2018.02068

15 Zeng, T. et al. Host- and stage-dependent secretome of the arbuscular mycorrhizal fungus Rhizophagus irregularis. Plant J 94, 411–425 (2018). 10.1111/tpj.13908

16 Wang, P. et al. A nuclear-targeted effector of Rhizophagus irregularis interferes with histone 2B mono-ubiquitination to promote arbuscular mycorrhisation. New Phytol 230, 1142–1155 (2021). 10.1111/nph.17236

17 De Mandal, S. & Jeon, J. Nuclear effectors in plant pathogenic fungi. Mycobiology 50, 259–268 (2022).

18 Tehrani, N. & Mitra, R. M. Plant pathogens and symbionts target the plant nucleus. Curr Opin Microbiol 72, 102284 (2023). 10.1016/j.mib.2023.102284

19 Harris, W., Kim, S., Vӧlz, R. & Lee, Y. H. Nuclear effectors of plant pathogens: Distinct strategies to be one step ahead. Mol Plant Pathol 24, 637–650 (2023). 10.1111/mpp.13315

20 Reddy, A. S., Marquez, Y., Kalyna, M. & Barta, A. Complexity of the alternative splicing landscape in plants. Plant Cell 25, 3657–3683 (2013). 10.1105/tpc.113.117523

21 Filichkin, S., Priest, H. D., Megraw, M. & Mockler, T. C. Alternative splicing in plants: directing traffic at the crossroads of adaptation and environmental stress. Curr Opin Plant Biol 24, 125–135 (2015). 10.1016/j.pbi.2015.02.008

22 Staiger, D. & Brown, J. W. Alternative splicing at the intersection of biological timing, development, and stress responses. Plant Cell 25, 3640–3656 (2013). 10.1105/tpc.113.113803

23 Staiger, D., Korneli, C., Lummer, M. & Navarro, L. Emerging role for RNA-based regulation in plant immunity. New Phytol 197, 394–404 (2013). 10.1111/nph.12022

24 Rigo, R., Bazin, J. R. M., Crespi, M. & Charon, C. L. Alternative Splicing in the Regulation of Plant-Microbe Interactions. Plant Cell Physiol 60, 1906–1916 (2019). 10.1093/pcp/pcz086

25 Laloum, T., Martin, G. & Duque, P. Alternative Splicing Control of Abiotic Stress Responses. Trends Plant Sci 23, 140–150 (2018). 10.1016/j.tplants.2017.09.019

26 Zhang, X. N. et al. Transcriptome analyses reveal SR45 to be a neutral splicing regulator and a suppressor of innate immunity in Arabidopsis thaliana. BMC Genomics 18, 772 (2017). 10.1186/s12864-017-4183-7

27 Xu, S. et al. Transportin-SR is required for proper splicing of resistance genes and plant immunity. PLoS Genet 7, e1002159 (2011). 10.1371/journal.pgen.1002159

28 Peng, S. et al. CONSTITUTIVE EXPRESSER OF PATHOGENESIS-RELATED GENES 5 is an RNA-binding protein controlling plant immunity via an RNA processing complex. Plant Cell 34, 1724–1744 (2022). 10.1093/plcell/koac037

29 Howard, B. E. et al. High-throughput RNA sequencing of pseudomonas-infected Arabidopsis reveals hidden transcriptome complexity and novel splice variants. PLoS One 8, e74183 (2013). 10.1371/journal.pone.0074183

30 Zheng, Y., Wang, Y., Ding, B. & Fei, Z. Comprehensive Transcriptome Analyses Reveal that Potato Spindle Tuber Viroid Triggers Genome-Wide Changes in Alternative Splicing, Inducible trans-Acting Activity of Phased Secondary Small Interfering RNAs, and Immune Responses. J Virol 91 (2017). 10.1128/JVI.00247-17

31 Huang, J. et al. Phytophthora Effectors Modulate Genome-wide Alternative Splicing of Host mRNAs to Reprogram Plant Immunity. Mol Plant 13, 1470–1484 (2020). 10.1016/j.molp.2020.07.007

32 Huang, J. et al. An oomycete plant pathogen reprograms host pre-mRNA splicing to subvert immunity. Nat Commun 8, 2051 (2017). 10.1038/s41467-017-02233-5

33 Jeong, B. R. et al. Structure function analysis of an ADP-ribosyltransferase type III effector and its RNA-binding target in plant immunity. J Biol Chem 286, 43272–43281 (2011). 10.1074/jbc.M111.290122

34 Mejias, J. et al. The root-knot nematode effector MiEFF18 interacts with the plant core spliceosomal protein SmD1 required for giant cell formation. New Phytol 229, 3408–3423 (2021). 10.1111/nph.17089

35 Jeong, S. SR Proteins: Binders, Regulators, and Connectors of RNA. Mol Cells 40, 1–9 (2017). 10.14348/molcells.2017.2319

36 Zhong, X. Y., Wang, P., Han, J., Rosenfeld, M. G. & Fu, X. D. SR proteins in vertical integration of gene expression from transcription to RNA processing to translation. Mol Cell 35, 1–10 (2009). 10.1016/j.molcel.2009.06.016

37 Ali, G. S. et al. Regulation of plant developmental processes by a novel splicing factor. PLoS One 2, e471 (2007). 10.1371/journal.pone.0000471

38 Zhang, X. N. & Mount, S. M. Two alternatively spliced isoforms of the Arabidopsis SR45 protein have distinct roles during normal plant development. Plant Physiol 150, 1450–1458 (2009). 10.1104/pp.109.138180

39 Carvalho, R. F. et al. The Arabidopsis SR45 Splicing Factor, a Negative Regulator of Sugar Signaling, Modulates SNF1-Related Protein Kinase 1 Stability. Plant Cell 28, 1910–1925 (2016). 10.1105/tpc.16.00301

40 Day, I. S. et al. Interactions of SR45, an SR-like protein, with spliceosomal proteins and an intronic sequence: insights into regulated splicing. Plant J 71, 936–947 (2012). 10.1111/j.1365-313X.2012.05042.x

41 Woodward, L. A., Mabin, J. W., Gangras, P. & Singh, G. The exon junction complex: a lifelong guardian of mRNA fate. Wiley Interdiscip Rev RNA 8 (2017). 10.1002/wrna.1411

42 Schlautmann, L. P. et al. Exon junction complex-associated multi-adapter RNPS1 nucleates splicing regulatory complexes to maintain transcriptome surveillance. Nucleic Acids Res 50, 5899–5918 (2022). 10.1093/nar/gkac428

43 Koroleva, O. A., Brown, J. W. & Shaw, P. J. Localization of eIF4A-III in the nucleolus and splicing speckles is an indicator of plant stress. Plant Signal Behav 4, 1148–1151 (2009).

44 Baldwin, K. L., Dinh, E. M., Hart, B. M. & Masson, P. H. CACTIN is an essential nuclear protein in Arabidopsis and may be associated with the eukaryotic spliceosome. Febs Lett 587, 873–879 (2013). 10.1016/j.febslet.2013.02.041

45 Venice, F. et al. At the nexus of three kingdoms: the genome of the mycorrhizal fungus Gigaspora margarita provides insights into plant, endobacterial and fungal interactions. Environ Microbiol 22, 122–141 (2020). 10.1111/1462-2920.14827

46 Morin, E. et al. Comparative genomics of Rhizophagus irregularis, R. cerebriforme, R. diaphanus and Gigaspora rosea highlights specific genetic features in Glomeromycotina. New Phytol 222, 1584–1598 (2019). 10.1111/nph.15687

47 Godfrey, D. et al. Powdery mildew fungal effector candidates share N-terminal Y/F/WxC-motif. BMC Genomics 11, 317 (2010). 10.1186/1471-2164-11-317

48 Stern, D. L. & Han, C. Gene Structure-Based Homology Search Identifies Highly Divergent Putative Effector Gene Family. Genome Biol Evol 14 (2022). 10.1093/gbe/evac069

49 Banani, S. F., Lee, H. O., Hyman, A. A. & Rosen, M. K. Biomolecular condensates: organizers of cellular biochemistry. Nat Rev Mol Cell Biol 18, 285–298 (2017). 10.1038/nrm.2017.7

50 Emenecker, R. J., Holehouse, A. S. & Strader, L. C. Biological Phase Separation and Biomolecular Condensates in Plants. Annu Rev Plant Biol 72, 17–46 (2021). 10.1146/annurev-arplant-081720-015238

51 Fu, X. D. & Maniatis, T. Isolation of a complementary DNA that encodes the mammalian splicing factor SC35. Science 256, 535–538 (1992). 10.1126/science.1373910

52 Zhang, Y. et al. Phase separation of HRLP regulates flowering time in Arabidopsis. Sci Adv 8, eabn5488 (2022). 10.1126/sciadv.abn5488

53 Ali, G. S., Golovkin, M. & Reddy, A. S. Nuclear localization and in vivo dynamics of a plant-specific serine/arginine-rich protein. Plant J 36, 883–893 (2003). 10.1046/j.1365-313x.2003.01932.x

54 Haynes, C. & Iakoucheva, L. M. Serine/arginine-rich splicing factors belong to a class of intrinsically disordered proteins. Nucleic Acids Res 34, 305–312 (2006). 10.1093/nar/gkj424

55 Cuevas-Velazquez, C. L. & Dinneny, J. R. Organization out of disorder: liquid-liquid phase separation in plants. Curr Opin Plant Biol 45, 68–74 (2018). 10.1016/j.pbi.2018.05.005

56 Ali, G. S., Prasad, K. V., Hanumappa, M. & Reddy, A. S. Analyses of in vivo interaction and mobility of two spliceosomal proteins using FRAP and BiFC. PLoS One 3, e1953 (2008). 10.1371/journal.pone.0001953

57 Golovkin, M. & Reddy, A. S. An SC35-like protein and a novel serine/arginine-rich protein interact with Arabidopsis U1-70K protein. J Biol Chem 274, 36428–36438 (1999). 10.1074/jbc.274.51.36428

58 Xing, D., Wang, Y., Hamilton, M., Ben-Hur, A. & Reddy, A. S. Transcriptome-Wide Identification of RNA Targets of Arabidopsis SERINE/ARGININE-RICH45 Uncovers the Unexpected Roles of This RNA Binding Protein in RNA Processing. Plant Cell 27, 3294–3308 (2015). 10.1105/tpc.15.00641

59 Malar, C. M. et al. Early branching arbuscular mycorrhizal fungus Paraglomus occultum carries a small and repeat-poor genome compared to relatives in the Glomeromycotina. Microb Genom 8 (2022). 10.1099/mgen.0.000810

60 Field, S., Jang, G. J., Dean, C., Strader, L. C. & Rhee, S. Y. Plants use molecular mechanisms mediated by biomolecular condensates to integrate environmental cues with development. Plant Cell 35, 3173–3186 (2023). 10.1093/plcell/koad062

61 Wang, W. & Gu, Y. The emerging role of biomolecular condensates in plant immunity. Plant Cell 34, 1568–1572 (2022). 10.1093/plcell/koab240

62 Huang, Y., Gattoni, R., Stevenin, J. & Steitz, J. A. SR splicing factors serve as adapter proteins for TAP-dependent mRNA export. Mol Cell 11, 837–843 (2003). 10.1016/s1097-2765(03)00089-3

63 Kumar, K. R. & Kirti, P. B. Novel role for a serine/arginine-rich splicing factor, AdRSZ21 in plant defense and HR-like cell death. Plant Mol Biol 80, 461–476 (2012). 10.1007/s11103-012-9960-8

64 Zhong, X. Y., Ding, J. H., Adams, J. A., Ghosh, G. & Fu, X. D. Regulation of SR protein phosphorylation and alternative splicing by modulating kinetic interactions of SRPK1 with molecular chaperones. Genes Dev 23, 482–495 (2009). 10.1101/gad.1752109

65 Du, J. et al. Nucleocytoplasmic trafficking is essential for BAK1- and BKK1-mediated cell-death control. Plant J 85, 520–531 (2016). 10.1111/tpj.13125

66 Pfaff, C. et al. ALY RNA-Binding Proteins Are Required for Nucleocytosolic mRNA Transport and Modulate Plant Growth and Development. Plant Physiol 177, 226–240 (2018). 10.1104/pp.18.00173

67 Tong, J. et al. ALBA proteins confer thermotolerance through stabilizing HSF messenger RNAs in cytoplasmic granules. Nat Plants 8, 778–791 (2022). 10.1038/s41477-022-01175-1

68 Chen, L. et al. Plant immunity suppressor SKRP encodes a novel RNA-binding protein that targets exon 3’ end of unspliced RNA. New Phytol (2023). 10.1111/nph.19236

69 Fu, Z. Q. et al. A type III effector ADP-ribosylates RNA-binding proteins and quells plant immunity. Nature 447, 284–288 (2007). 10.1038/nature05737

70 Streitner, C. et al. An hnRNP-like RNA-binding protein affects alternative splicing by in vivo interaction with transcripts in Arabidopsis thaliana. Nucleic Acids Res 40, 11240–11255 (2012). 10.1093/nar/gks873

71 Nicaise, V. et al. Pseudomonas HopU1 modulates plant immune receptor levels by blocking the interaction of their mRNAs with GRP7. EMBO J 32, 701–712 (2013). 10.1038/emboj.2013.15

72 Zorin, E. A. et al. Transcriptome Analysis of Alternative Splicing Events Induced by Arbuscular Mycorrhizal Fungi (Rhizophagus irregularis) in Pea (Pisum sativum L.) Roots. Plants (Basel) 9 (2020). 10.3390/plants9121700

73 Almagro Armenteros, J. J., et al. SignalP 5.0 improves signal peptide predictions using deep neural networks. Nat Biotechnol 37, 420–423 (2019). 10.1038/s41587-019-0036-z

74 Horton, P. et al. WoLF PSORT: protein localization predictor. Nucleic Acids Res 35, W585–587 (2007). 10.1093/nar/gkm259

75 Ishida, T. & Kinoshita, K. PrDOS: prediction of disordered protein regions from amino acid sequence. Nucleic Acids Res 35, W460–464 (2007). 10.1093/nar/gkm363

76 Erdos, G., Pajkos, M. & Dosztanyi, Z. IUPred3: prediction of protein disorder enhanced with unambiguous experimental annotation and visualization of evolutionary conservation. Nucleic Acids Res 49, W297–W303 (2021). 10.1093/nar/gkab408

77 Sievers, F. et al. Fast, scalable generation of high-quality protein multiple sequence alignments using Clustal Omega. Mol Syst Biol 7, 539 (2011). 10.1038/msb.2011.75

78 Robert, X. & Gouet, P. Deciphering key features in protein structures with the new ENDscript server. Nucleic Acids Res 42, W320–324 (2014). 10.1093/nar/gku316

79 Crooks, G. E., Hon, G., Chandonia, J. M. & Brenner, S. E. WebLogo: a sequence logo generator. Genome Res 14, 1188–1190 (2004). 10.1101/gr.849004

80 Liu, W. et al. IBS: an illustrator for the presentation and visualization of biological sequences. Bioinformatics 31, 3359–3361 (2015). 10.1093/bioinformatics/btv362

81 Tamura, K., Stecher, G. & Kumar, S. MEGA11: Molecular Evolutionary Genetics Analysis Version 11. Mol Biol Evol 38, 3022–3027 (2021). 10.1093/molbev/msab120

82 Letunic, I. & Bork, P. Interactive Tree Of Life (iTOL) v5: an online tool for phylogenetic tree display and annotation. Nucleic Acids Res 49, W293–W296 (2021). 10.1093/nar/gkab301

83. Lueck, S. & Douchkov, D. “Blaster” - a graphical user interface for common sequence analysis tools. Journal of Open Source Software 4, 10.21105/joss.01411 (2019).

84 Raudvere, U. et al. g:Profiler: a web server for functional enrichment analysis and conversions of gene lists (2019 update). Nucleic Acids Res 47, W191–W198 (2019). 10.1093/nar/gkz369

85 Kuhn, H., Kuster, H. & Requena, N. Membrane steroid-binding protein 1 induced by a diffusible fungal signal is critical for mycorrhization in Medicago truncatula. New Phytol 185, 716–733 (2010). 10.1111/j.1469-8137.2009.03116.x

86 Heck, C. et al. Symbiotic Fungi Control Plant Root Cortex Development through the Novel GRAS Transcription Factor MIG1. Curr Biol 26, 2770–2778 (2016). 10.1016/j.cub.2016.07.059

87 Karimi, M., Inze, D. & Depicker, A. GATEWAY vectors for Agrobacterium-mediated plant transformation. Trends Plant Sci 7, 193–195 (2002). S1360-1385(02)02251-3 [pii]

88 Limpens, E. et al. Formation of organelle-like N2-fixing symbiosomes in legume root nodules is controlled by DMI2. Proceedings of the National Academy of Sciences of the United States of America 102, 10375–10380 (2005).

89 Schutze, K., Harter, K. & Chaban, C. Bimolecular fluorescence complementation (BiFC) to study protein-protein interactions in living plant cells. Methods Mol Biol 479, 189–202 (2009). 10.1007/978-1-59745-289-2_12

90 Pham, G. M. et al. Construction of a chromosome-scale long-read reference genome assembly for potato. Gigascience 9 (2020). 10.1093/gigascience/giaa100

91 Kim, D., Langmead, B. & Salzberg, S. L. HISAT: a fast spliced aligner with low memory requirements. Nat Methods 12, 357–360 (2015). 10.1038/nmeth.3317

92 Pertea, M. et al. StringTie enables improved reconstruction of a transcriptome from RNA-seq reads. Nat Biotechnol 33, 290–295 (2015). 10.1038/nbt.3122

93 Trapnell, C. et al. Differential gene and transcript expression analysis of RNA-seq experiments with TopHat and Cufflinks. Nat Protoc 7, 562–578 (2012). 10.1038/nprot.2012.016

94 Kong, L. et al. CPC: assess the protein-coding potential of transcripts using sequence features and support vector machine. Nucleic Acids Res 35, W345–349 (2007). 10.1093/nar/gkm391

95 Langmead, B. & Salzberg, S. L. Fast gapped-read alignment with Bowtie 2. Nat Methods 9, 357–359 (2012). 10.1038/nmeth.1923

96 Li, B. & Dewey, C. N. RSEM: accurate transcript quantification from RNA-Seq data with or without a reference genome. BMC Bioinformatics 12, 323 (2011). 10.1186/1471-2105-12-323

97 Love, M. I., Huber, W. & Anders, S. Moderated estimation of fold change and dispersion for RNA-seq data with DESeq2. Genome Biol 15, 550 (2014). 10.1186/s13059-014-0550-8

98 Shen, S. et al. rMATS: robust and flexible detection of differential alternative splicing from replicate RNA-Seq data. Proc Natl Acad Sci U S A 111, E5593–5601 (2014). 10.1073/pnas.1419161111

99 Gietz, R. D. & Woods, R. A. Transformation of yeast by lithium acetate/single-stranded carrier DNA/polyethylene glycol method. Methods Enzymol 350, 87–96 (2002).

100 Voinnet, O., Rivas, S., Mestre, P. & Baulcombe, D. An enhanced transient expression system in plants based on suppression of gene silencing by the p19 protein of tomato bushy stunt virus. Plant J 33, 949–956 (2003).

101 Rech, S. S., Heidt, S. & Requena, N. A tandem Kunitz protease inhibitor (KPI106)-serine carboxypeptidase (SCP1) controls mycorrhiza establishment and arbuscule development in Medicago truncatula. Plant J 75, 711–725 (2013). 10.1111/tpj.12242

102 Win, J., Kamoun, S. & Jones, A. M. Purification of effector-target protein complexes via transient expression in Nicotiana benthamiana. Methods Mol Biol 712, 181–194 (2011). 10.1007/978-1-61737-998-7_15

103. Schindelin, J., et al. Fiji: an open-source platform for biological-image analysis. Nat Methods 9, 676–682 (2012). 10.1038/nmeth.2019

104 RStudio Team. RStudio: Integrated Development Environment for R. . RStudio, PBC, Boston, MA, URL http://www.rstudio.com/ (2021).

## References

Haynes C, Iakoucheva LM. 2006. Serine/arginine-rich splicing factors belong to a class of intrinsically disordered proteins. Nucleic Acids Res 34(1): 305–312.

## References

Wang BB, Brendel V. Molecular characterization and phylogeny of U2AF35 homologs in plants. Plant Physiol. 2006 Feb;140(2):624–36. doi: 10.1104/pp.105.073858.

Golovkin M, Reddy AS. Structure and expression of a plant U1 snRNP 70K gene: alternative splicing of U1 snRNP 70K pre-mRNAs produces two different transcripts. Plant Cell. 1996 Aug;8(8):1421–35. doi: 10.1105/tpc.8.8.1421. PMID: 8776903; PMCID: PMC161266.

Golovkin M, Reddy AS. An SC35-like protein and a novel serine/arginine-rich protein interact with Arabidopsis U1-70K protein. J Biol Chem. 1999 Dec 17;274(51):36428–38. doi: 10.1074/jbc.274.51.36428. PMID: 10593939.

Tanabe N, Kimura A, Yoshimura K, Shigeoka S. Plant-specific SR-related protein atSR45a interacts with spliceosomal proteins in plant nucleus. Plant Molecular Biology. 2009 Jun;70(3):241–252. DOI: 10.1007/s11103-009-9469-y.

